# Two stochastic processes shape diverse senescence patterns in a single-cell organism

**DOI:** 10.1101/105387

**Authors:** Ulrich K. Steiner, Adam Lenart, Ming Ni, Peipei Chen, Xiaohu Song, François Taddei, Ariel B. Lindner, James W. Vaupel

**Author notes:** Current address: BGI Shenzhen, Shenzhen, China. Current address: National Center for Nanoscience and Technology, Beijing, China (Peipei Chen). Joint senior authors (Ariel B. Lindner, James W. Vaupel). Equal contribution of second authors (Adam Lenart, Ming Ni).

## Abstract

Despite advances in aging research, a multitude of aging models, and empirical evidence for diverse senescence patterns, understanding is lacking of the biological processes that shape senescence, both for simple and complex organisms. We show that for a isogenic *Escherichia coli* bacterial population senescence results from two stochastic processes. A primary random deterioration process within the cell, such as generated by random accumulation of damage, leads to an exponential increase in mortality early in life followed by a late age mortality plateau; a secondary process of stochastic asymmetric transmission of an unknown factor at cell fission influences mortality. This second process is required to explain the difference between the classical mortality plateaus detected for young mothers’ offspring and the near non-senescence of old mothers’ offspring as well as the lack of a mother offspring correlation in age at death. We observed that life span is predominantly determined by underlying stochastic stage dynamics. Our findings support models based on stage-specific actions of alleles for the evolution of senescence. This support might be surprising since these models that have not specifically been developed in the context of simple, single cell organisms. We call for exploration of similar stochastic influences beyond simple organisms.

## Introduction

One of the major challenges for biodemographic research on aging is to understand what drives senescence patterns (Vaupel et al. 1998, López-Otín et al. 2013). This challenge is illustrated by the variety of aging models that assume different, as yet unverified generating processes (Hamilton 1966, Kirkwood 2005, Wachter et al. 2014). Prominent — mutually not exclusive — evolutionary theories of senescence, such as William’s antagonistic pleiotropy hypothesis (Williams 1957), or Medawar’s mutation accumulation hypothesis (Medawar 1952), provide general predictions about uniform senescence patterns across many taxa (Hamilton 1966). However, these generalities have been questioned both theoretically and empirically by illustrating how negligible and negative-senescence can theoretically be achieved and been empirically found in various species (Vaupel et al. 2004, Jones et al. 2014). Despite an extensive literature on mechanistic approaches of aging, the generating processes that drive such diversity in senescence patterns remain opaque (López-Otín et al. 2013). Mechanistic approaches to aging identify a multitude of underlying biochemical, molecular and organismal mechanisms that relate to the decline of function with age, many of which are rooted in direct or indirect oxidative processes (Kirkwood 2005, López-Otín et al. 2013). Examples include age-related mitochondrial dysfunction, telomere shortening, stem cell exhaustion, genotypic instability, epigenetic alterations, accumulation of damaged proteins and general loss of proteostasis (Lindner and Demarez 2009, Tyedmers et al. 2010, López-Otín et al. 2013). Yet, researchers have not conclusively determined whether such mechanisms are a cause or consequence of aging. This failure may be due to the complexity of model systems of aging. As a consequence, it is difficult to relate these mechanisms directly to the observed demographic patterns (Tyedmers et al. 2010, López-Otín et al. 2013, Denoth Lippuner et al. 2014). Only such linkage — between mechanisms and senescence patterns — can elucidate generating processes that underlie the various theories and aspects of aging.

Aging in bacteria has been established over the last one and a half decades and thereby provided a simple biological system to study aging (Ackermann et al. 2003, Stewart et al. 2005). Before, bacteria have been thought to not age (Williams 1957), because they normally fission into two equal sized progeny. These progeny were assumed to be identical, that is, the original mother cell would die when fissioning and leaves two identical daughters (Johnson and Mangel 2006, Tyedmers et al. 2010). This perspective has changed because the resulting progeny are phenotypically unequal (Tyedmers et al. 2010). This asymmetry among the progeny is manifested by asymmetry of intracellular content at cell fission and between carrying old and newly formed cellular poles (Stewart et al. 2005, Lindner et al. 2008). Here, we follow the convention of old pole and new pole cells, being referred to as mother and daughter cell respectively (Fig. 1), to track the age of individual cells (Stewart et al. 2005). It has been shown that mother cells have a higher probability to accumulate misfolded protein and grow slower as compared to their daughter cells, but a causal relationship to mortality was not established (Lindner et al. 2008). From a theoretical point of view asymmetry is required to rejuvenate some cells in order to prevent whole population aging. Otherwisepopulations would accumulate more and more damage if perfect symmetric fissions occurred and damage accumulation accedes damage repair and dilution due to growth and fission (Ackermann et al. 2007, Evans and Steinsaltz 2007).

Here, we reveal characteristics of the underlying aging processes by inference from observed senescence patterns. We achieve this aim by quantifying demographic parameters of a simple biological system, isogenic individual *E. coli* bacteria cells, under highly controlled environmental conditions. We used a high-throughput microfluidic device (Fig. 1; movie S1) to track individual cells throughout their lives (Wang et al. 2010, Gasset-Rosa et al. 2014, Jouvet et al. 2017). Two types of cells were tracked: early daughter cells and late daughter cells (Fig. 1). A late daughter cell is the last daughter cell produced by an early daughter and hence an offspring of an old mother. An early daughter cell is the offspring of a young mother since they were haphazardly extracted from a population that grew exponentially. According to stable stage population theories such exponentially growing populations are vastly dominated by young cells (Caswell 2001)(Fig. S1). We expect early daughters to hold little or no deterioration or damage at birth, whereas late daughters are likely be born with some damage. In this study, we use damage as a synonym for any unknown aging factor that leads to deterioration and increased mortality. We use this synonym because many aging factors are assumed to be accumulated damage caused by oxidative processes (Kirkwood et al. 2005, López-Otín et al. 2013). A third group of cells, resembling the late daughter type are the last daughter cells produced by the late daughters, which we call second generation late daughters (Results only in SI1). Our definition of mother and daughter cells builds on the concept of cell polarity for both early daughter and late daughter cells to track the age of cells and distinguish between the mothers (old pole cell) and the daughters (new pole cell) (Fig.1).

## Material and Methods

For brevity we only provide in this section an overview of the methods. For more detailed methodological information on strains and growth conditions, time-lapse imaging, image analysis, determining death and estimating demographic parameters, statistical analyses, the simulations please see the supplemental information.

We collected data in two independent sets of experiments. We loaded *E. coli* K12-derived MG1655 strain cells into a designed microfluidic (PDMS) chip (Wang et al. 2010, Gasset-Rosa et al. 2014, Jouvet et al. 2017) from an exponentially growing culture in supplemented minimum media M9. During each experiment, we acquired 77 hours of time lapse phase-contrast imaging (15 frames/hour for each of the 2×44 fields followed) using a temperature-controlled inverted microscope, at 43°C with an accuracy of the temperature control at the chip of ±0.1°C (note within the chip temperature should be even more closely controlled due to some buffering of the chip). We used 43°C to accelerate the aging process and thereby shorten the time the system needed to run under stable conditions. Such stability was particularly important to accurately track late daughter and second generation late daughter cells. Increasing temperature up to 43 °C scales senescence patterns, but does not alter the shape of patterns (Jouvet & Steiner unpublished). At 44 °C patterns got unstable and cells were not viable over longer time periods (results not presented). The rod-shaped bacteria cells grew in dead end side channels of a microfluidic chip, with the focal cells trapped at the end of the side channels, and we tracked them over their lifespan (Fig. 1; movie S1). We used customized image analysis to generate the demographic data (lifespan, cell elongation rate, cell size, and time of each division).

We assured by starting the experiments with exponentially growing cells that the initial loaded cells are descendants of young mothers (i.e. they are early daughter cells) (Fig. S1). At the end of their lives, these early daughter cells produced a last daughter (late daughters) that then became the next bottom-most cell trapped at the end of the respective growth channel. Therefore, we could directly compare mother and daughter cells. Note that the late daughters are not born at the same time (main text Fig. 2 E, F). The late daughter cells produced another generation of late daughter cells at the end of their lives (second generation late daughter cells) for which the results are shown in Fig. S2-8, Table S1-3. Analysis on the empirical data were done in program R (R Core Team 2016) using general linear, generalized linear, and non-linear models. Models were selected based on information criteria (AIC) (Burnham and Anderson 1998) or based on differential evolution algorithm for global optimization (R package DEoptim).

Simulations and extending random deterioration models.

For the extended random deterioration model, we first estimated parameters by fitting a fixed frailty model — a Gamma-Gompertz-Makeham (GGM) model — to the observed survival data of the early daughter cells (Yashin et al. 1994, Missov and Vaupel 2015). We then translated these fitted GGM model parameters to an extended random deterioration process model, an extended LeBras type deterioration model (Le Bras 1976). In doing so, we took advantage of mathematical similarities between the two types of models, even though they are biologically distinct (Yashin et al. 1994, Missov and Vaupel 2015). With these random deterioration model parameters we could estimate the probability matrix of an individual being at stage *i* at age *x*. Microsimulations based on these models provided stage at death distribution. For the late daughters, we assumed that they are born at a scaled version of the same damage stage in which their mothers died. We also assumed that late daughters accumulate damage at the same rates as early daughters do, i.e. same probabilities to transition to a higher damage state of early and late daughters. We further assumed that the amount of accumulated damage in late daughters had the same effect on mortality than on early daughters, that is, early and late daughter cells are not fundamentally different except that late daughters are born in different damage stages, while early daughters start their lives without damage, but the baseline mortality (Makeham term) can be interpreted as some starting level of damage even for early daughters. Our model simplifies the biological system substantially in as much as no repair of damage or purging of damage through asymmetric division at cell fission is considered. Damage accumulates unidirectionally, mortality increases exponentially with accumulated damage, and each cell suffers from an age-independent baseline mortality risk (Makeham term).

**Fig. 1:**
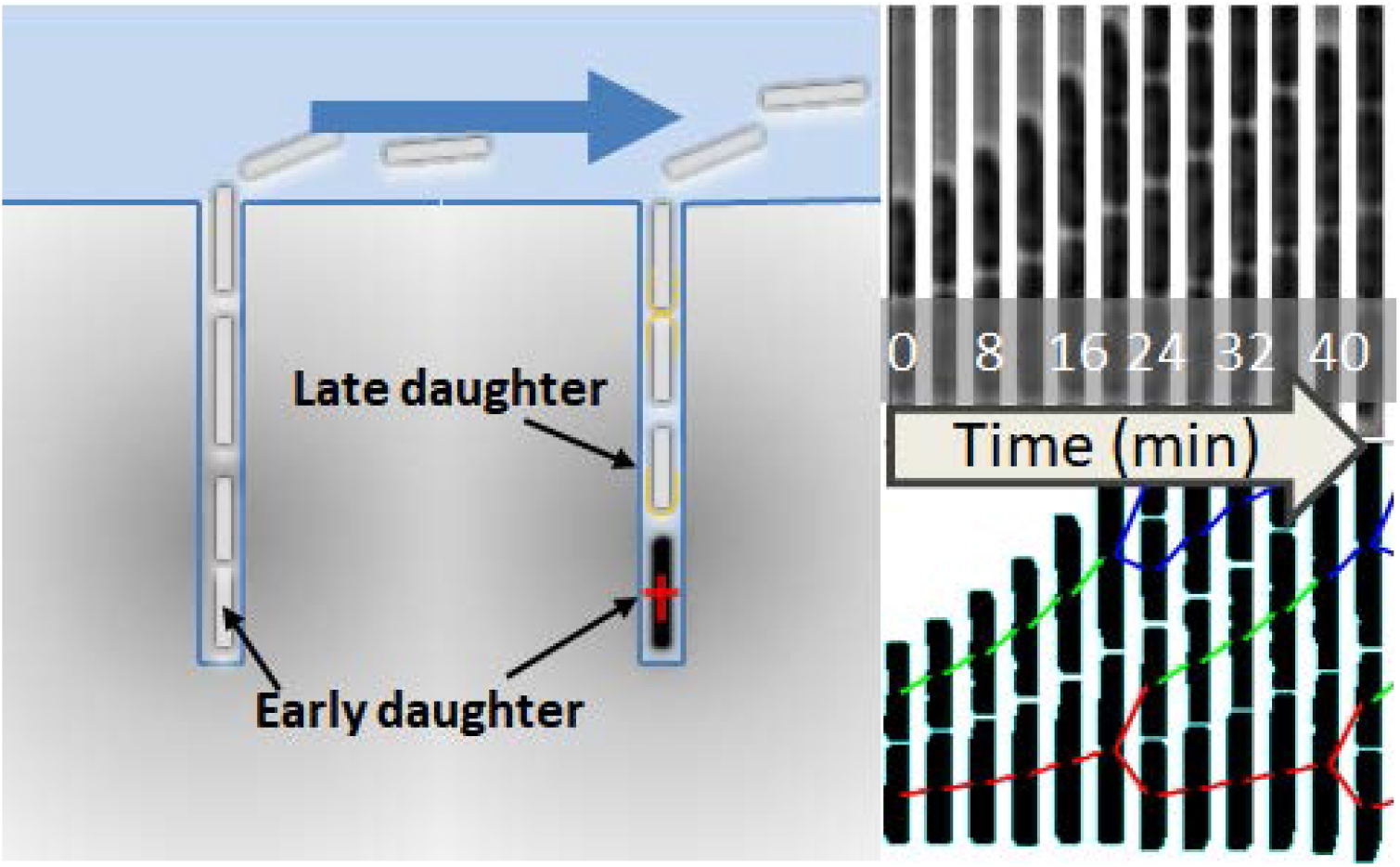
Left panel: Overview sketch of the microfluidic device with dead-end side channels and the main laminar flow channel where the media is flushed through. The early daughters (founding initial loaded mothers) are the bottommost cells of the dead-end channels. Their daughters (new pole progenitor cells) are located closer to the main laminar flow channel and have more recent poles. When the mother (early daughter) dies (side channel on the right), its last daughter (1^st^ generation late daughter) is then tracked throughout its life. Accordingly, 2^nd^ generation late daughter cells are tracked once their mother (1^st^ generation late daughter) dies. Right top and bottom panel: Phase contrast time sliced (4 min intervals) side channel images with initial loaded early daughter at the bottom (old pole progenitor). Growth (cell elongation) and division can be tracked throughout their life as depicted in the segmented cell lineages (bottom right panel). See also movie S1.

## Results and Discussion

### Chronological aging in *E. coli*

In our experiments, we followed by automated time-lapse microscopy 516 early daughters throughout their lives as they divide and age in microfluidic dead-end channels (Fig. 1). We also followed two generations of late daughters of these 516 early daughters and recorded each cells growth rate and lifespan. In this, we reveal classical senescence patterns of a decrease in reproduction (Fig. 2A, B) — indicated by decreased cell elongation and increased size at division with age (Fig. S2, Table S1) — for both early and late daughters (Figure 2A, B). The observed senescence patterns (Fig. 2C, D) describe chronological aging in *E. coli* and support previous studies on replicative senescence (time counted as number of divisions) in this species (Lindner et al. 2008, Wang et al. 2010) (Fig. S2 & Fig. S4). The main result highlights that early daughters and late daughters differ fundamentally in their senescence pattern, even though they are isogenic and grown in a highly controlled environment. Only the early daughters exhibit the classical senescence pattern, marked by an early exponential increase of the probability of death followed by a later age mortality plateau. Late daughters have the same probability of death across most of their lives, i.e. no senescence is observed at the population level. Only late in life does mortality increase, but this increase is largely driven by only one data point, the one for the last age class (>30h) and is accompanied by increased uncertainty due to small sample size at that age. Such a plateau, exhibited by the early daughters — recurrent in many higher organisms including humans (Vaupel et al. 1998) — has not been previously shown for bacteria, potentially indicating deep-rooted features of aging and senescence. In yeast, ambiguous results on senescence patterns have been described (Minois et al. 2005, Denoth Lippuner et al. 2014).

**Fig. 2:**
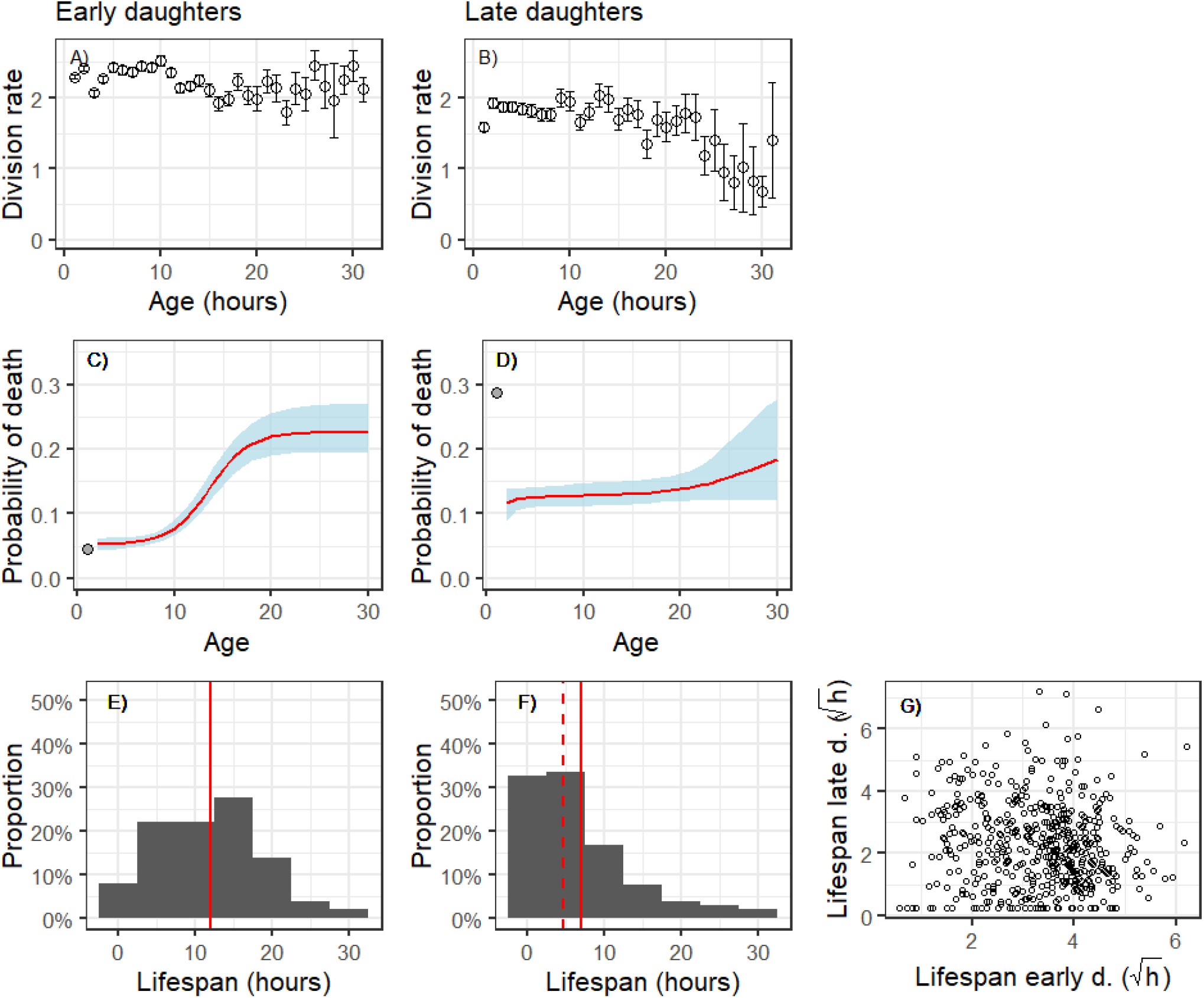
Division rate (divisions per hour) (A, B) and probability of death per hour (C, D) with increasing age, as well as lifespan distribution (in hours) (E, F) of isogenic *E. coli* cells grown under highly controlled environmental conditions in a microfluidic device (Fig. 1, movie S1). Age patterns are shown for early daughters (A, C, E; N=516), and late daughters (B, D, F; N=516). Late daughters are the last offspring of the founding early daughters. The correlation of early daughter’s lifespan (mothers’ lifespan) (square root transformed for better visibility; N=516) versus the lifespan of their last daughter cell (late daughter cells; N=516) is shown in panel (G). For (A, B) hourly means ± standard errors are plotted, for (C, D) the fitted regression ± 95% confidence intervals are plotted (confidence intervals are estimated based on individual level data SI1). The fitted regression (red lines) are likelihood optimization for Gamma-Gompertz-Makeham functions and relate closely to the modeling approach we took below (C, D, see also Fig. 3, simulations below, and SI1; Table S1, Table S2 for statistical testing) (Burnham and Anderson 1998). The grey points in panel C, D, mark average first hour probability of death which have been excluded from the modeling. For lifespan distributions (E, F), mean and median are marked by solid and dashed lines respectively. For patterns of the second generation late daughters, see Table S2, Table S3, Table S4, and Fig. S3, Fig. S7, Fig. S8, Fig. S9, Fig. S10, Fig. S11.

The probability of death of newborn late daughters (<=1hour, filled grey data point in Fig. 2D) drops after the first hour to a level that is lower than their long-lived mothers (early daughters above age 20) and then remains at that level throughout their lives. This drop in mortality early in life of late daughters suggests a damage purging effect (see below). Such a drop of mortality after birth might indicate heterogeneity among newborns amenable to evolutionary selective forces (Yashin et al. 1994), e.g. heterogeneity in maternally-transmitted damage between mothers and daughters. With increasing age we detected increased variance and increased uncertainty of parameter estimation (Fig. 2 A-E). Such increase is expected for age-structured demographic analyses and well understood for its declining number of individuals with age (Brillinger 1986, Klein and Moeschberger 2003, Scherbov and Ediev 2011). The increase in variance is more pronounced for probabilities of death (Fig. 2 C, D) than for division rates (Fig. 2 A, B), because each individual dies only once while each individual divides multiple times throughout its life and hence division rates suffer less from smaller sample sizes. Binning over larger time spans or having an earlier open age bracket would diminish the effect of increased variance with age.

### Damage accumulation and selection

If we assume, as in most physiological theories of aging, that accumulated damage is the determining factor for the probability of death, then the observed constant probability of death with age for older early daughters and the late daughters (Fig. 2C, D) indicates an equilibrium distribution of damage among individuals. Such equilibrium is realized at the population level while the individuals accumulate damage in a stochastic manner. It equilibrates on one hand the accumulation of new damage within individuals similar to processes described by random deterioration models (Weitz and Fraser 2001), and on the other hand, the intracellular repair of damage and purging of damage by two mechanisms. The first mechanism reduces damage within an individual by asymmetric division of damage at cell fission, which increases variance in damage among individuals (mother and daughters). The second mechanism selects against highly damaged cells, i.e. the most damaged cells of a population die, which lowers the average level of damage in the surviving population, and reduces the variance of damage among individuals (Evans and Steinsaltz 2007). Intriguingly, this equilibrium (plateau level) is lower for late daughters (Fig. 2D, see Table S3) than the late-age mortality plateau of the early daughters (Fig. 2C). This indicates — based on fixed frailty models (Missov and Vaupel 2015) — higher heterogeneity in transmitted damage of late daughters compared to early daughters. Such increased heterogeneity at birth predicts an earlier onset of selection against highly damaged cells which would explain why in our simulations (below) late daughters have on average less damage compared to early daughters at the plateau (see Fig. 3A, B, Fig. S10). Purging of damage through asymmetric division at cell fission has been shown in yeast, where late daughters showed reduced lifespan, but through further division and presumably damage dilution, the descendants of these late daughter yeast cells recovered high longevity (Kennedy 1994). This is also reminiscent of asymmetric partition of protein aggregates in bacterial cells due to their passive accumulation in bacterial mother cells (Lindner et al. 2008, Coquel et al. 2013).

### Stochastic variability in lifespan and lack of mother-daughter correlation of lifespan

We did not find any influence of the lifespan of the mothers on the lifespan of their (last) daughters (Fig. 2G) (Table S6, Fig. S8). Since we are working with an isoclonal population we do not expect any genetic variability among mother-daughter pairs; therefore, we would not expect a genetically driven mother-daughter correlation of lifespan caused by additive genetic differences. However, non-genetic factors such as maternal transmitted damage would predict mother-daughter correlated lifespans except if i) daughters would be perfectly rejuvenated — all daughters would be born without damage (see also Fig. 3E) —, or ii) if the transmission of damage to the daughter is independent of the amount of accumulated damage by the mother — the mother-daughter transmission of damage is stochastic. Since late daughters display higher probability of death from birth onwards (Fig. 2D), which is indicative of some maternally transmitted damage and obviously shows that late daughters are not perfectly rejuvenated, we can exclude the first explanation why a mother-daughter correlation of lifespan is lacking. This leaves the second explanation where the amount of damage transmitted from the mother to the daughter is stochastic and independent of the age of the mother. Early daughters seem to hold little or no damage at birth, since their mothers have not yet accumulated much damage, whereas the average late daughter seems to be born with some damage that raises its probability of death. The lack of a mother-daughter correlation of lifespan is also remarkable as it contrasts heritability of lifespan in humans and other complex organisms that show similar senescence patterns (Finch and Kirkwood 2000).

The lack of correlation between mother-daughter lifespan suggests that the assumed mother-daughter transmission of damage is predominantly stochastic. If damage accumulates stochastically within a cell over its life, long-lived mothers might have been lucky and accumulated damage at a lower rate than the average mother, but they still accumulate damage with increasing age. At age of death, long-lived mothers might have similar damage levels compared to short lived mothers. Short-lived mothers, which likely accumulated damage at an exceptional high rate, do not produce daughters with shorter life expectancy. Despite their difference in lifespan, neither long-lived nor short-lived mothers tend to transmit higher or lower amounts of damage to their daughters, otherwise we would find a negative or positive correlation in mother-daughter lifespan (Fig. 2G). Therefore, our observation that early daughters — which are assumed to hold little or no damage at birth — die at very different ages indicates that the process of damage accumulation within an individual appears to be stochastic and varies highly among individuals. Despite an isogenic population in a highly controlled homogeneous environment, we observe a high variation in lifespan and reproduction among individuals, both for early daughters (Median±stdev lifespan 12 ± 7 hours, Coefficient of Variation, CV 0.57, Fig. 2E) (Fig. S7) and late daughters (Median±stdev 5 ± 7; CV=1.01, Fig. 2F). Such high variation supports substantial influences of stochastic events in shaping lifespan.

### Modeling damage accumulation and asymmetric transmission of damage

To support our conclusions based on the empirical results and to gain a better understanding of the underlying stochastic processes that shape our observed senescence patterns, we extended random deterioration process models (e.g. random accumulation of damage, see details in Methods and SI1) (Le Bras 1976, Yashin et al. 1994, Evans and Steinsaltz 2007). Our extended model assumes that all cells accumulate damage unidirectionally at the same rate, i.e. cells move with equal probability to a higher damage state, mortality increases exponentially with accumulated damage, and each cell suffers from an age-independent baseline mortality risk. We also assume that early daughters are born without damage (Fig. 3A). The results of our simulations show that with increasing age population level damage (and variance) slowly increases at early ages, accelerates after 10 hours and plateaus after 20 hours (Fig. 3A). These patterns in damage accumulation mirror the observed mortality patterns (Fig. 2C). Mortality plateaus, as observed for early daughters, can be explained by either fixed frailty, i.e. heterogeneity in damage stage at birth, or acquired heterogeneity throughout life (Vaupel and Yashin 1985, Yashin et al. 1994, Weitz and Fraser 2001, Missov and Vaupel 2015). In our system heterogeneity in damage stage at birth seems to have little influence, at least for early daughter cells. Senescence is driven by an age independent baseline mortality and accrued damage as in random deterioration models (Le Bras 1976, Yashin et al. 1994, Weitz and Fraser 2001). This conclusion is supported by two results of our model. First, most early daughters die with no or very little damage (Fig. S10), and second, the observed survival patterns follow closely the survival curve of simulated cells that did not accumulate any damage throughout life (Fig. 3C).

Contrasting the assumption for the early daughter cells, in our simulation the average simulated late daughter is born with some damage (~24% are born without damage) (Yashin et al. 1994)(Fig. 3B). For the following results, it is important to note that this heterogeneity in damage at birth of late daughters is an underestimation since we do not include the exceptionally high first-hour mortality rate (grey filled data point in Fig. 2D) for the simulations. In our simulations the distribution of this damage at birth of the late daughters is related to the distribution of the accumulated damage at death of the early daughter cells (mothers of the late daughters, Fig. S10). For the simulations, we assumed that the mother to daughter transmission of damage is a fixed fraction of all the damage the mother accumulated over her lifespan. Note, we only simulated a single binomial division, rather than multiple asymmetric fissions as others have done (Evans and Steinsaltz 2007). We computed the fixed fraction of damage transmission between mother and daughter by optimizing the survival pattern predicted by the simulations to the observed survival curves.

We found that the optimal (simulated) fixed fraction was at a low level of 0.07. This indicates a highly asymmetric division of damage between mothers and daughters. If damage transmission between mothers and daughters would be symmetrical (0.5), the predicted survival of the simulations (Fig. 3D hatched line) does not match the observed survival (Fig. 3D red line). For early daughters, the level of asymmetry makes little difference because most of them die with no or little accumulated damage (Fig. 3C, Fig. S10). The predicted survival pattern of late daughters at older ages slowly converges to the simulated survival patterns of undamaged cells (outer right grey line) (Fig. 3D). Therefore, many cells in our simulation that survive to old ages hold little damage, which supports our interpretation of our experimental data that late daughter cells are born with diverse damage states, highly damaged cells are selected against, and at old ages, population-level survival patterns are strongly influenced by cells that have not accumulated much damage.

Similar selective effects have been described by population level demographic theories as heterogeneity’s ruses, where selection among different damage stages (fixed frailty; i.e. heterogeneity in damage stage at birth) leads to diverse senescence patterns at the population level (Vaupel and Yashin 1985). Compared to our simple model that only considers a single binary fission (branching) and suggests a strong asymmetry, more complex and realistic models that include multiple branching (fission) events show less extreme asymmetry at cell fission to be adaptive (Evans and Steinsaltz 2007). This difference in optimal asymmetry between our simple and the more complex models can be understood in that the asymmetric transmission of damage is distributed over multiple fissions in the more complex models, while for our model it is concentrated at a single event. Even though the optimized fixed transmission fraction between mother and daughters of our model is low at 0.07, such a fixed fraction would lead to a correlation in lifespan between mother and daughters of about 0.25 (Fig. 3E). If different levels of fixed transmissions are simulated, correlation in lifespan between mother and daughters increases with increasing transmission. Only when there is no transmission of any damage (perfect rejuvenation of the daughters) we see no correlation in lifespan (Fig. 3E). The observed data does not support such correlation in lifespan (Fig. 2G), but also does not support perfect rejuvenation (Fig. 2C and D). Hence these simulations support our conclusion that the fraction of transmitted accumulated damage from mother to daughter at cell fission varies stochastically and is not fixed at a low level of 0.07 as our model assumes.

**Fig. 3:**
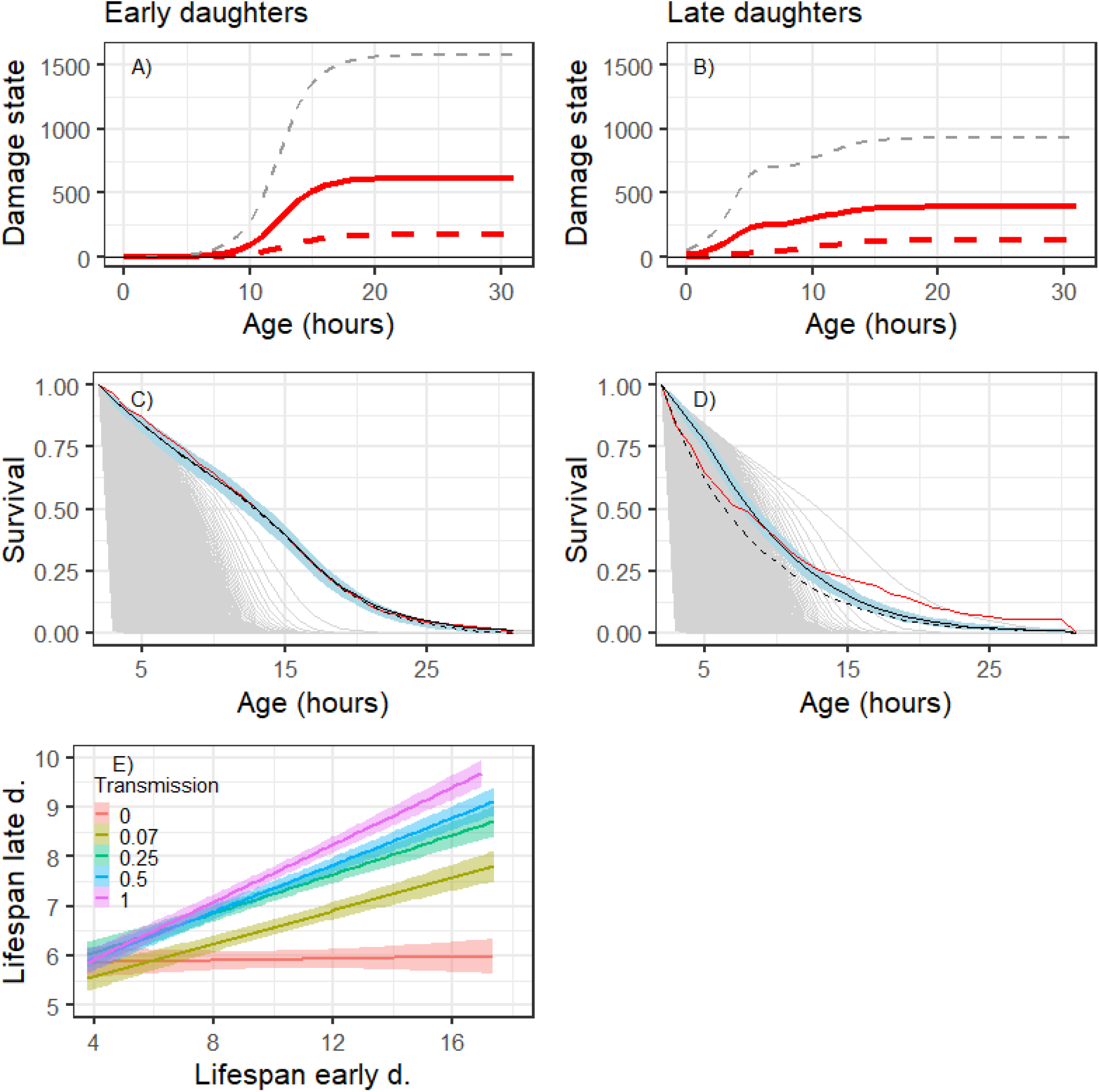
Random deterioration model: Simulation results are shown of mean (red solid line, + SD grey hatched lines) and median (red hatched line) damage state with increasing age (A, B); observed experimental population level survival curves (red solid line; see also related probability of death Fig. 2 C, D and related observed distributions of death Fig. 2E, F), Gamma-Gompertz-Makeham simulated survival curves with symmetric damage transmission (hatched black line) as well as simulated asymmetric damage transmission of 7% (solid black lines ± 95% CI in blue) (C, D). Graphs are shown for early daughter cells (A, C) and late daughter cells (B, D). Thin grey lines in (C, D) depict expected survival curve of simulated cells with different fixed damage state: outermost (right) curve depicts the highest survival of simulated cells with no damage throughout their lives, most left survival curve depicts the lowest survival of simulated cells that were born with maximum damage level of 5000. Lifespan (square root transformed) of simulated 516 early daughter cells (mothers) versus the lifespan of their simulated last daughter cell (late daughter cells) with different levels of mother to daughter damage transmission (E). The different scenarios include the optimized fixed transmission level at 0.07 (forest green), a scenario for perfect rejuventation, i.e. 0 transmission (red), 0.25 transmission (green), 0.5 transmission, i.e. symmetric (equal) transmission between mother and daughter (blue), and transmission of all accumulated damage to the daughter (1) (pink).CI are shown for each correlation in lifespan between mothers and daughters as shaded areas.

In this study, our experimental setup limits our analysis to two extreme cases: early and late daughter cells (in fact the last daughter cells). We hypothesize that the results do not only hold for these two extremes but rather portray the extremes of a continuous process across different aged cells. Two observations support our claim: first, changes in mortality of early daughters (Fig. 2C) and age patterns in reproduction (Fig. 2A) are somewhat gradual (more so for survival than for reproduction), suggesting a gradual underlying mechanism; second, we do not observe a pronounced pattern just before death (Fig. S5 & Fig. S6, Table S5). Such a pattern would be expected if the last daughters are exceptional because their mothers are approaching death, for instance in the way as predicted under terminal investment theories (Clutton-Brock 1984).

### Two stochastic processes shape diverse senescence patterns

We conclude that the diverse senescence patterns, including the classical senescence pattern of a mortality plateau, are determined by two stochastic processes that relate to underlying (damage) states and only indirectly to age (Yashin et al. 1994, Wachter 1999, Weitz and Fraser 2001, Evans and Steinsaltz 2007). The primary process is a random deterioration process, e.g. the stochastic accumulation of damage throughout life, and the secondary process involves the stochastic transmission of damage from the mother to their daughters at cell fission. This transmitted damage or some other unknown aging factor increases the probability of death but is “non-heritable” as we show by the lack of correlation between mother and daughter lifespan (Yashin et al. 1994, Wachter 1999, Weitz and Fraser 2001, Evans and Steinsaltz 2007). Not only additive genetic variation but also other commonly assumed drivers of senescence, such as epigenetic variability, age itself, or the (extrinsic) environment can be ruled out as major players in our study. If epigenetic variability had a significant influence, a positive correlation among mother and daughter lifespan would be seen. If chronological age determined senescence, early and late daughter cells would show similar mortality patterns, hence senescence is rather driven by stage dynamics than by chronological age. The highly controlled microfluidic system creates a uniform environment and hence we can exclude extrinsic environmental drivers; this does not imply that senescence patterns do not differ under different environmental conditions, we just investigated only one specific environment. For instance, under complex media variance in division size and related division rate is increased compared to the minimum medium we used (Gangan and Athale 2017). Minimum medium also decreases the rate of filamentation, a stress response that causes continued cell elongate without dividing. Recovery of filamentous cells is rarely observed under minimum medium but frequent under complex medium (Wang et al. 2010) (see also SI). The near identical conditions cells experience in the microfluidic system decreases environmental variability compared to naturally experienced micro-habitats. The laminar flow provides large amounts of fresh media that constantly diffuses within seconds into the dead-end side channels and therefore prohibits built-up of micro-environmental-niches. Though, such micro-environments (e.g. biofilms) are characteristic for the natural conditions in the guts of the mammalian hosts most *E. coli* life and have evolved under. The microfluidic system we use largely prevents cell-to-cell contact and chemical communication among cells that can influence population growth (Aoki et al. 2005). The highly controlled conditions therefore lowers phenotypic variability compared to natural conditions. To what degree senescence patterns are altered by the environment remains to be explored. Temperature seem to conserve the shape of senescence patterns and simply scales patterns differently, whereas nutrient concentrations and nutrient source might alter shapes as has been found in other organisms (Stroustrup et al. 2016)(Steiner unpublished).

Our findings indicate that two simple stochastic processes can create complex senescence patterns. Easily understood, simplistic, and plausible arguments behind evolutionary theories of aging might lure readers into the misconception that chronological age in itself plays an important role in senescence. Such simplistic assumptions of age-specific drivers must be approached with caution and seem not applicable to this study. Selective forces likely decline with time in our system, but the decline might be dominated by underlying stage dynamics. Selection on heterogeneity among individuals is rather generated by such stage dynamics than by increasing chronological age. In this sense, selective forces and their decline vary considerably among individuals, even though genetic load that relates to the accumulation of deleterious mutations should be negligible in an isoclonal population as ours. The substantial effect of stochastic events on life histories is also indicated by the large variability in life histories despite excluding any genetic or environmental variability. Such large stochastic variation supports arguments behind neutral theories of life history (Steiner and Tuljapurkar 2012) that suggest that stochastic variability can account for substantial fractions of the total variability in fitness components. Our interpretation of stochastic events determining individual life courses is consistent with findings of significant stochastic influences at the molecular (Elowitz et al. 2002) and protein level (Tyedmers et al. 2010, Balázsi et al. 2011) and adds to the growing interest in such phenotypic heterogeneity and individuality (Wolf et al. 2007, Davidson and Surette 2008, Ackermann 2015).

Future challenges will include determining whether these stochastic processes are neutral or adaptive, and what drives their evolution (Steinsaltz and Evans 2004, Kærn et al. 2005, Norman et al. 2015, Vera et al. 2016). For the basic aging processes that drive senescence, the causal relationships that drive age patterns are currently unknown (Lindner and Demarez 2009, López-Otín et al. 2013). The complex senescence patterns found in this study of a simple model organism under highly controlled conditions emphasizes the challenges to quantify contributions of well-defined determinants of aging in the complex systems on which most aging research is focused (López-Otín et al. 2013). Comparing mean characteristics of differently aged individuals as frequently done in aging research might hold limited insights in system where determining stages are likely highly dynamic. In light of the growing evidence that stochastic processes can have cascading effects across all levels of higher organisms (Finch and Kirkwood 2000, Elowitz et al. 2002, Balázsi et al. 2011) new avenues in aging research may require a shift towards the underlying stochastic processes that drive such stage dynamics in simple systems like bacteria and perhaps beyond. Identifying the underlying currently unknown (damage) states remains another challenge that we believe requires a combined quantitative demographic and mechanistic approach, because of the high level of stochastic influences. Promising steps in such directions have been initiated by theoretically exploring stage-specific alleles shaping senescence patterns (Wachter et al. 2014) and by exploring transcription factor signal dynamics at the single cell level across increasing parts of the lifespans of many individuals (Norman et al. 2013).

## Acknowledgements

We thank D.A. Roach, D. Steinsaltz, T. Coulson, Y. Yang and S. Tuljapurkar for discussions and comments.

## Supporting information SI1

### Strains and growth conditions

We fabricated the microfluidics chips as previously described (Gasset-Rosa et al. 2014). We grew *E. coli* K12 MG1655 (Blattner 1997) strain derivative (intC∷pSulA-yfp) cells at 43°C in filtered minimal medium supplemented with 0.4% Glucose and 0.2% Casamino Acids (hereafter M9). We diluted 100μl of the overnight culture into 50ml of M9 and then grew it at 43°C to exponential growth phase (OD600 ~0.2). We centrifuged the cells and resuspended the cell pellet in 100μl of M9 to load them into the chip by injection into the main (feeding) channel followed by centrifugation (15 minutes at ~168g). We applied a continuous laminar flow (2.7ml/h) of M9 supplemented with 1.5% Polyethylene Glycol (PEG P3015 Sigma Aldrich) through the main channel throughout the experiment. The temperature was kept constant at 43°C (see below).

### Time-lapse imaging

We followed bacterial growth in the microfluidics chip by phase-contrast time-lapse imaging at a rate of 4min/frame using a MetaMorph (Molecular Devices)-controlled inverted Nikon microscope (Eclipse Ti, 100xobjective, CoolSNAP HQ2 camera) with a temperature-control chamber (Live Imaging Services). We continuously scanned 44 positions, each comprising 18 channels, for 77 hours, in two independent experimental sets.

### Image analysis

We used a custom image analysis to segment all cells within the side channels per frame, the software measured the cells’ location and size within the time series and generated cell lineages. To this end, phase contrast time-lapse images were lowpass-filtered and background flattened (MetaMorph Molecular Devices software) to increase contrast. We further processed the images by a customized ImageJ plugin for cell segmentation to crop each of the growth channels, stabilize the time sequenced images, adjust the brightness and to create a binary image by thresholding and filtering (median filter). Finally, the ImageJ plugin water-sheds the time sliced images. We then applied a final round of segmentation in Matlab by correcting errors in segmentation by checking for minimum (0.8) and maximum (1.4) cell elongation rates for each cell (size time-4 min, size time+4min). This way, we recorded for any time points (at 4 minute intervals) the exact location of a cell in the side channel (the pixel coordinate of the center of mass of the cell area), the side channel of the cell, and if a cell divided. By the position within the side channel we determined which cell was the mother (old pole progenitor) cell (bottommost cell), their most recent daughter (new pole progenitor) cell and so forth. We measured the length of a cell as the largest distance of the rod shaped cell in the orientation of the side channel. We applied a minimum “cell length” of ~1μm to exclude artefacts originating mainly from small deformations in the growth channels, or some shadows/reflections generated at the dead end of the growth channel.

### Age at death

We defined age at death as the time when a cell stopped growing and dividing for at least one hour and 20 min. None of the cells divided after such a long division arrest or resumed growth.

### Early and late daughter cells

We defined the early daughter population as the cells that were initially loaded into the microfluidic chip and settled at the dead-end of the side channels. They originated largely from young mothers because the population was in exponential growth phase when we loaded cells into the chip (Jagers 1978). The age distribution of these early daughter cells is, because of the exponential growth phase, negatively exponentially distributed. We show their expected age distribution in Fig. S1, based on a Leslie matrix generated from the survival and division rates of the early daughters (Fig. 2A & C). Due to the exponential growth phase, we assume a stable age distribution (the right eigenvector corresponding to the dominant eigenvalue of the Leslie matrix) (Caswell 2001). We tracked these early daughter cells to the end of their lives (Fig. 1, movie S1). The last daughters that were produced by these early daughter cells are the late daughter population. These late daughters are all of age zero but originate from mother cells (the early daughters) that had accumulated damage over their lives, and that were only one division from dying. We again tracked these late daughters throughout their lives. We focus mainly on these first two types of cells (early and late daughters), though as expected, the next generation of late daughter cells (second generation late daughters) follow the late daughter patterns (Fig. S4-6).

**Fig. S1:**
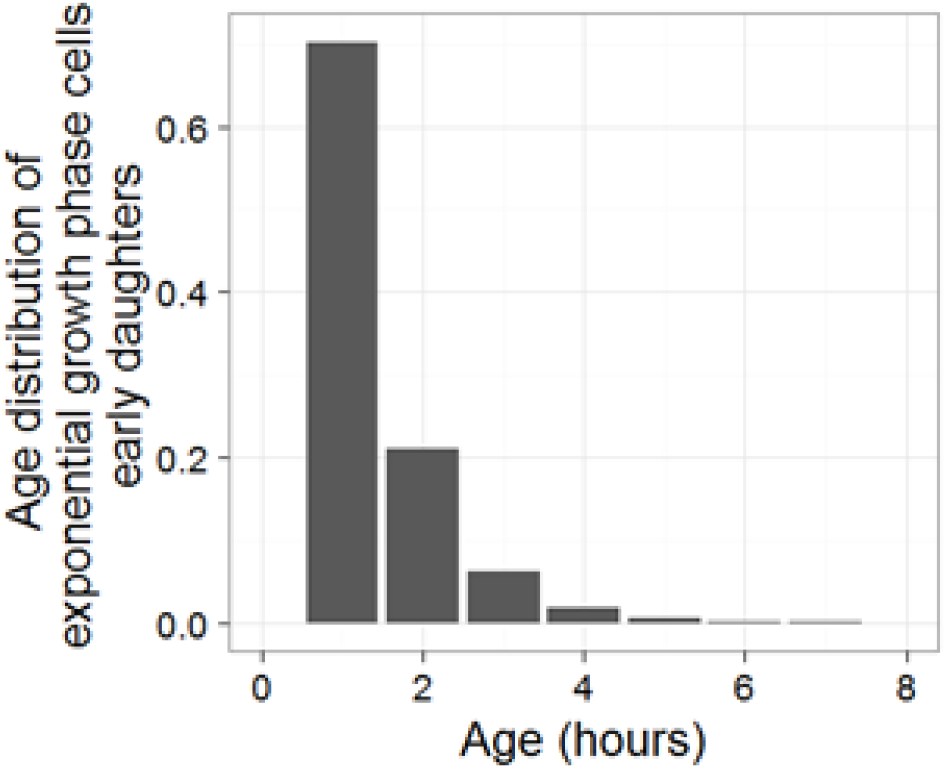
Expected (stable) age distribution (in hours) of early daughters (initial loaded cells), estimated from a Leslie matrix parameterized with the demographic rates of the early daughter cells. Due to the theory of stable age populations, the distribution of the ages of the mothers of these early daughters should be exactly as the (stable) age distribution shown here of the initial loaded (early daughter cells) at the start of the experiments.

### Estimating demographic parameters

Image analysis (see above) provided us the basis for estimating the age-specific demographic parameters: division rate, cell elongation rate, size at division, and survival rate. We excluded all cells that were not tracked throughout their lives. This concerned mostly cells that filamented (cell division arrested without growth arrest) and were flushed out of the side channels when they reached a size longer than 25 μm (length of the side channel). We also excluded all (early and late daughter) cells that died at chronological age 0 (never started to grow or divide). This was practiced because loading into the chip might be damaging and we wanted to exclude such death. This approach was conservative, because fewer early daughters never started growing or divided compared to late daughters (who could not have been damaged by the loading). We used data from 516 early daughters that produced 516 late daughters which in turns produced 298 (second generation) late daughters that were tracked throughout their lives.

### Statistical analysis

We did the statistical analysis using the R software (R Core Team 2016). We estimated and plotted hourly rates for division rates, cell elongation, and cell size at division rather than rates on the 4 min intervals because data would be sparse for 4 min intervals at older ages. Already the hourly rates at old ages suffer from sparse estimates and increased uncertainty. We also clustered all data of ages beyond 30 h for the same reason. To guide statistical testing, we used model selection (Burnham and Anderson 1998) based on AIC (Akaike Information Criteria) and considered — as common practice —better support between models when the ΔAIC was more than 2. For the parameter estimation of the GGM and extended random deterioration models, we used the DEoptim R package and the optim function within the stats R package. We calculated the CI (Fig. 2 C & D) by bootstrapping 1000 times. This bootstrapping is based on individual level data not the hourly means Fig. 2 C, D. We choose such an likelihood based global optimization approach to closely relate to the GGM modeling approach we used for the simulations. Using model comparison (AIC) based on the hourly rates (results not shown) selects for models with sigmoidal shapes like the GGM likelihood optimization, i.e. an early exponential increase in mortality followed by a late age mortality plateau (Fig. 2 C). For early daughters, other exponential or linear functions receive less support. For late daughters model comparisons selected non-senescence (Fig. 2 D, flat mortality) over linear or exponential models. We did not test for any breakpoint models or step functions, since such models are not expected based on evolutionary theories of aging.

### Lifespan distribution second generation late daughters

In the main article, we focused on early and late daughter cells. Qualitatively, patterns of the second gen eration late daughters (the last daughters produced by the late daughters) correspond to those of their m others (the late daughters), though data becomes sparse uncertainty increases and some deviations occur. Fig. S7 illustrates lifespan distributions of the 298 second generation late daughters, their mothers (late daughters), and grandmothers (early daughters). Kolmogorov-Smirnov tests (two-sided) verifies that both lifespan distributions between early and late daughters (D = 0.3691, p < 0.00001), and early to second generation late daughters are significantly different (D = 0.4094, p < 0.00001), while those between late daughters and second generation late daughters do not differ (D = 0.0973, p = 0.1189).

**Fig. S7:**
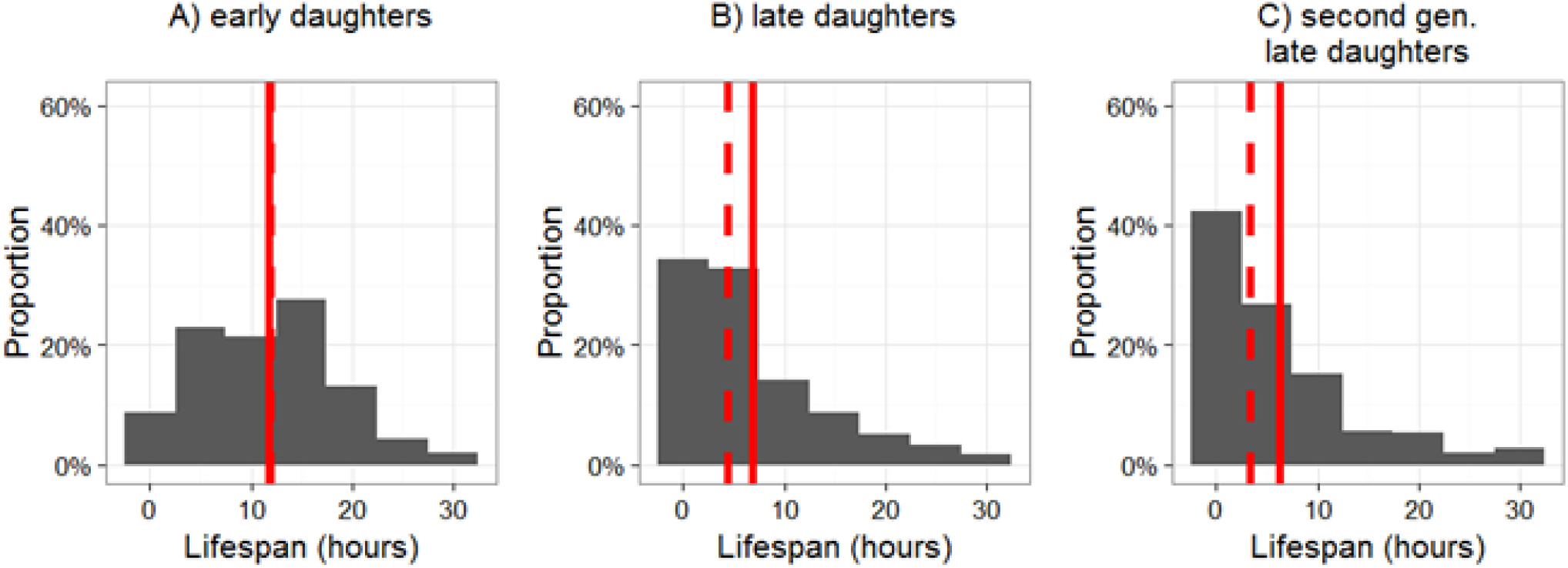
Lifespan distribution (in hours) of isogenic *E. coli* cells grown under identical environmental conditions in a microfluidic device (Fig. S1). 1A – Founding early daughter cells (Median±stdev lifespan 12 ± 6.8 hours, Coefficient of Variation, CV 0.57) (N=298); 1B – Late daughters (last daughter) of the founding early daughters of 1A (Median±stdev lifespan 4.4 ± 7.0 hours, Coefficient of Variation, CV 1.03) (N=298); 1C – Second generation late daughters (last daughter of the late daughters of 1B; (Median±stdev lifespan 3.4 ± 7.1 hours, Coefficient of Variation, CV 1.14) (N=298). (Fig. S1 & S3). Hence, early daughters are the mothers of late daughters and grandmothers of second generation late daughters.

### Heritability or cross generation correlation in lifespan

In order to estimate correlation of lifespan between mother cells (early daughters) and their last (late) daughter cells, we used linear models on square root transformed data for age at death (lifespan) for the mothers’ lifespan as response variable and their last daughters’ lifespan as explanatory variable, or intercept only models (null model) as comparative models.

Model selection (Table S6) does not allow distinguishing between the two alternative models and we ca nnot completely rule out that the longer-lived mothers produce slightly shorter-lived late daughters (Fig. 2G). In any case, the effect size is weak and such a negative correlation would suggest that longer-live d mothers accumulate more damage that is then partly transmitted to their late daughter. Similar results hold for late daughters to second generation late daughters (Fig. S8). We cannot completely rule out a w eak tendency that long-lived (late daughter) mothers produce last daughters (second generation late dau ghters) that live slightly shorter (Table S6). In any case, our simulations with a low fixed transmission r ate of 7% of damage transmitted between mother and daughters would lead to much higher correlation i n lifespan as observed (Fig. 3E main text).

**Fig. S8:**
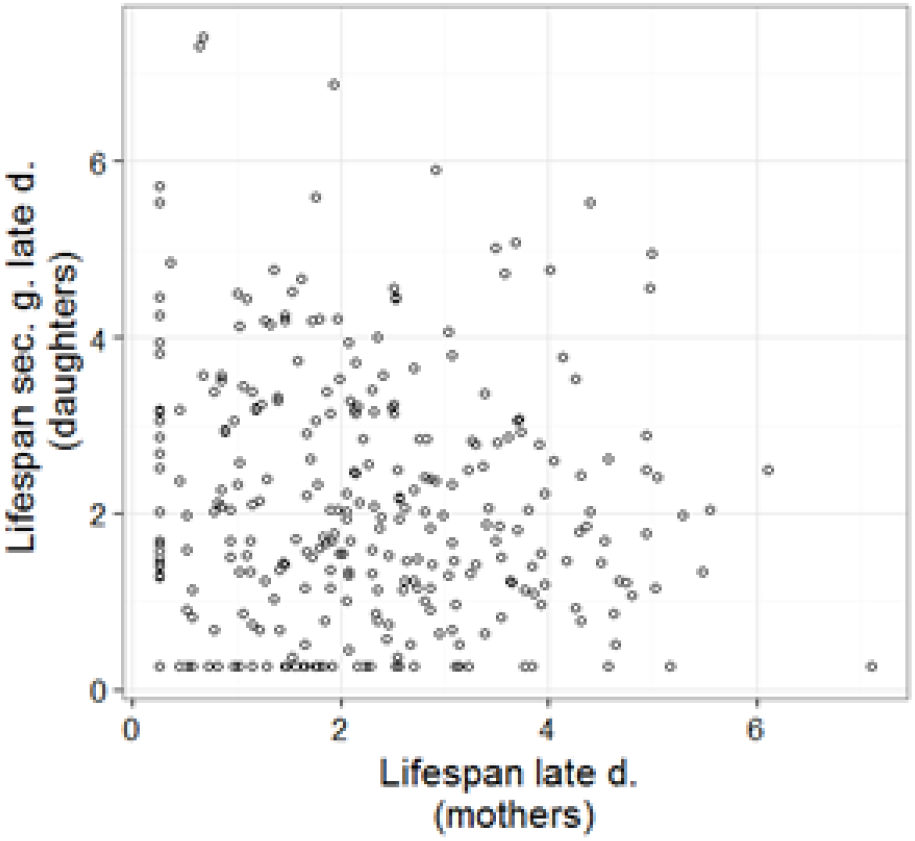
Correlation between lifespan (Square root-transformed) of the 298 late daughters [mothers] versus the lifespan of their last daughter cell, the second generation late daughters (Table S6).

**Table S6:** Model selection for correlation between lifespan (Square root transformed, Gaussian error)

| Model | Slope | Std. Error | T value | p-value | AIC | ΔAIC |
| --- | --- | --- | --- | --- | --- | --- |
| Early daughters [516 mothers] and daughter cell (last daughter of early daughter) |  |  |  |  |  |  |
| sqrt(Lifespan Daughters) ~ sqrt(Lifespan mothers) | -0.10 | 0.05 | -1.8 | 0.07 | 1774.1 |  |
| sqrt(Lifespan Daughters) ~ intercept only |  |  |  |  | 1775.5 | 1.3 |
| **Late daughters [mothers 298] and second generation late daughters (last daughter cell produced by late daughters)** |  |  |  |  |  |  |
| sqrt(Lifespan Daughters) ~ sqrt(Lifespan mothers) | -0.09 | 0.06 | -1.55 | 0.12 | 1061.1 |  |
| sqrt(Lifespan Daughters) ~ intercept only |  |  |  |  | 1061.5 | 0.4 |

### Lifespan and lifetime reproductive success: reproductive and chronological aging

Previous aging studies on *E. coli* (Stewart et al. 2005, Wang et al. 2010) and other bacteria (Ackermann et al. 2003) or many yeast studies (Denoth Lippuner et al. 2014) have focused on replicative aging and patterns have been described for time measured in number of divisions. Such age measures can be difficult to compare directly to chronological age if the time between divisions varies. Nonetheless, we detected a strong correlation between lifespan and the lifetime reproductive success (number of divisions an individual undergoes throughout its life), but there remains some variation, particularly for longer lived individuals (Fig. S4).

**Fig. S4:**
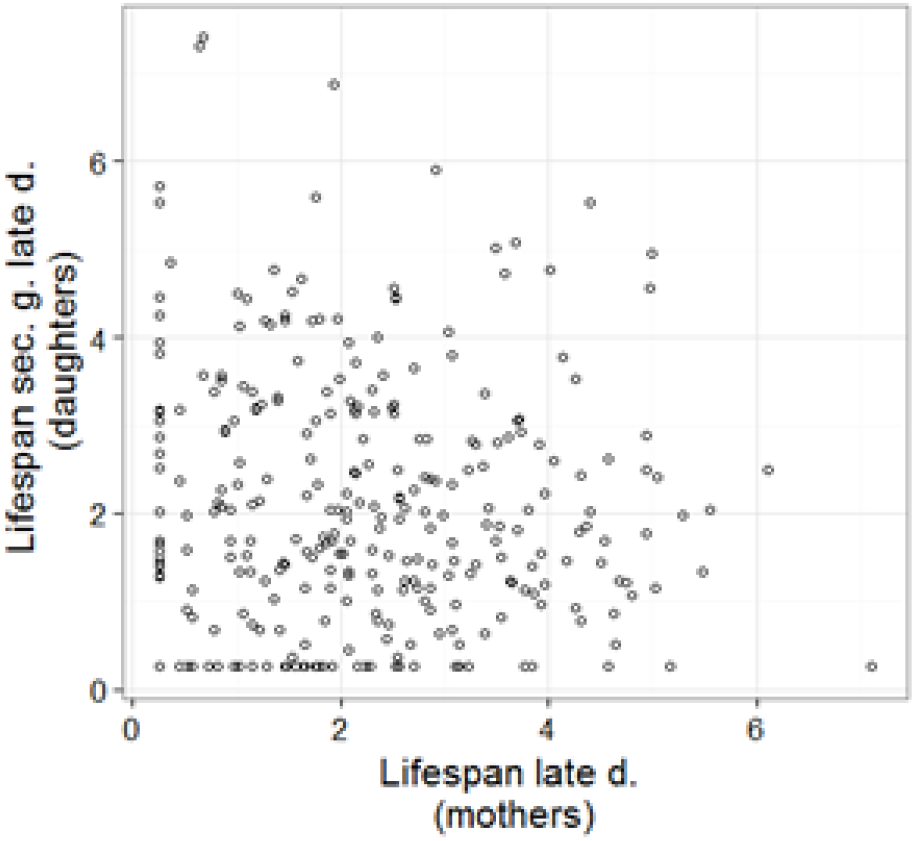
Correlation between lifespan and the lifetime reproductive success (cumulative number of divisions an individual undergoes throughout its life) for left panel) the 516 early daughters, mid panel) the 516 late daughters, and right panel) the 298 second generation late daughters.

### Division rate

We fitted linear models on hourly average division rates (at the cellular level) as response variable and age as explanatory variable. A model with a linear and quadratic term was best supported for the early (Fig. 2A), late (Fig. 2B) and second generation late (Fig. S3A) daughters (Table S2).

**Table S2:** Model selection division rate

| Model | Slope±Std. Error | Quadratic<br>term±StdE | AIC | ΔAIC |
| --- | --- | --- | --- | --- |
| **Early daughters** |  |  |  |  |
| Divison rate ~intercept<br>only |  |  | 19663 | 24 |
| Divison rate ~age | -0.011±0.002 |  | 19642 | 21 |
| **Divison rate**<br>**~age+age^2** | **0.005±0.0078** | **-0.0007±0.0003** | **19638** |  |
| **Late daughters** |  |  |  |  |
| Divison rate ~intercept<br>only |  |  | 12432 | 24 |
| Divison rate ~age | -0.009±0.004 |  | 12428 | 20 |
| **Divison rate**<br>**~age+age^2** | **0.039±0.011** | **-0.002±0.0005** | **12408** |  |
| **Second generation late daughters** |  |  |  |  |
| Divison rate ~intercept<br>only |  |  | 5494 | 63.3 |
| Divison rate ~age | 0.009±0.004 |  | 5490.3 | 59.6 |
| **Divison rate**<br>**~age+age^2** | **0.090±0.011** | **-0.004±0.0004** | **5430.7** |  |

### Mortality rate

We fitted Gamma-Gompertz-Makeham related models to the survival data. GGM models were tested for assumptions among parameters, for instance, whether shape and scale parameters are independent or defined by a rate parameter. For late (Fig. 2D) and second generation late daughters (Fig. S3B), we optimized the scaling of the mother to daughter state relationship (transmission factor) based on the age at death state distribution of early daughters (or late daughters for second generation late daughters) (Fig. S10). We assumed that accumulation of damage, which is the stage transition probabilities did not differ among early, late and second generation late daughters. We also assumed that the effect of the stage was not different for early, late or second-generation late daughters. These assumptions allowed us to use the translated GGM estimates of the early daughters also for late and second generation late daughters.

**Fig. S3:**
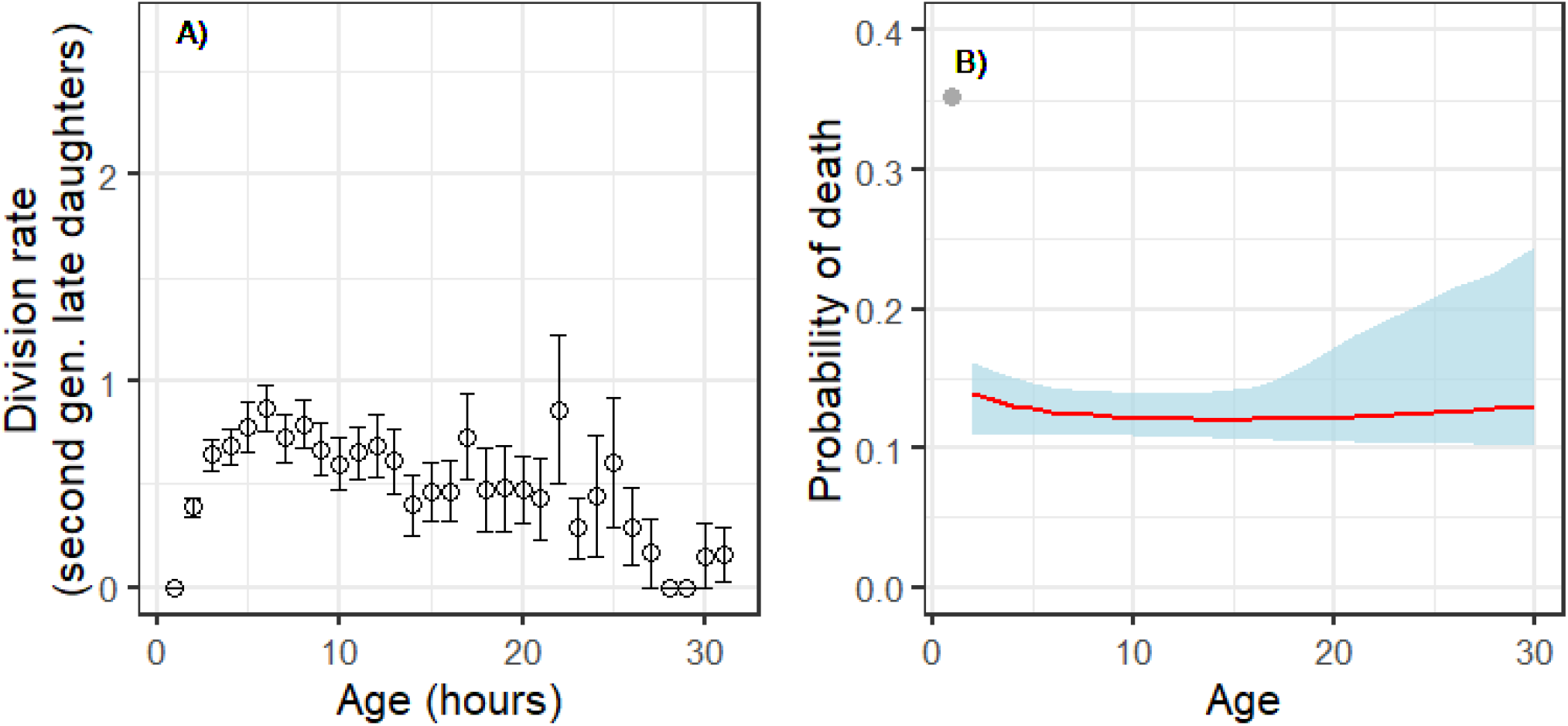
Age-specific division rate (number of divisions per hour) (A) and age-specific hourly mortality rate (B) of the 298 second generation late daughters. For (A) means ± standard errors are plotted, for (B) fitted regression ± 95% Confidence Intervals are plotted. Associated model selection results are presented in Table S2.

### Difference in mortality rates at old ages for early and late daughters

The late age mortality plateau detected in early daughters (Fig. 2C) and the flat mortality hazard pattern of the late daughters might indicate similar mortality rates at older ages (>19 hours). To test for this we compared models that distinguished between early and late daughters but only included cells that survived to an age of at least 20 hours. Late daughter cells have lower mortality rates (qx=0.1248±0.0191 mean±Std.Err.) than early daughters at old ages (qx=0.2005±0.0168 mean±Std.Err.) (Table S3). This result comprises 73 early daughters and 41 late daughters that lived longer than 19 hours.

**Table S3:** Model selection for age-specific mortality rates (qx) of old (>19h) early daughters and old (>19h) late daughters, based on binomial models.

| Model | Estimate±Std. Error | Quadratic term | AIC | ΔAIC |
| --- | --- | --- | --- | --- |
| Mortality ~intercept only |  |  | 95.79 | 6.09 |
| **Mortality ~cell type** | **-0.591±0.211** |  | **89.7** |  |

### Cell elongation rates

We illustrate findings for cell elongation rates for early, late and second generation late daughters (Fig. S2). When comparing linear models, the best fitted model for all three cell types was a model with a linear and quadratic term, though the curvature was minimal for early daughters (Table S1). Early and late daughters showing a decline in cell elongation rate with age, while second generation late daughters patterns are less clear.

**Fig. S2:**
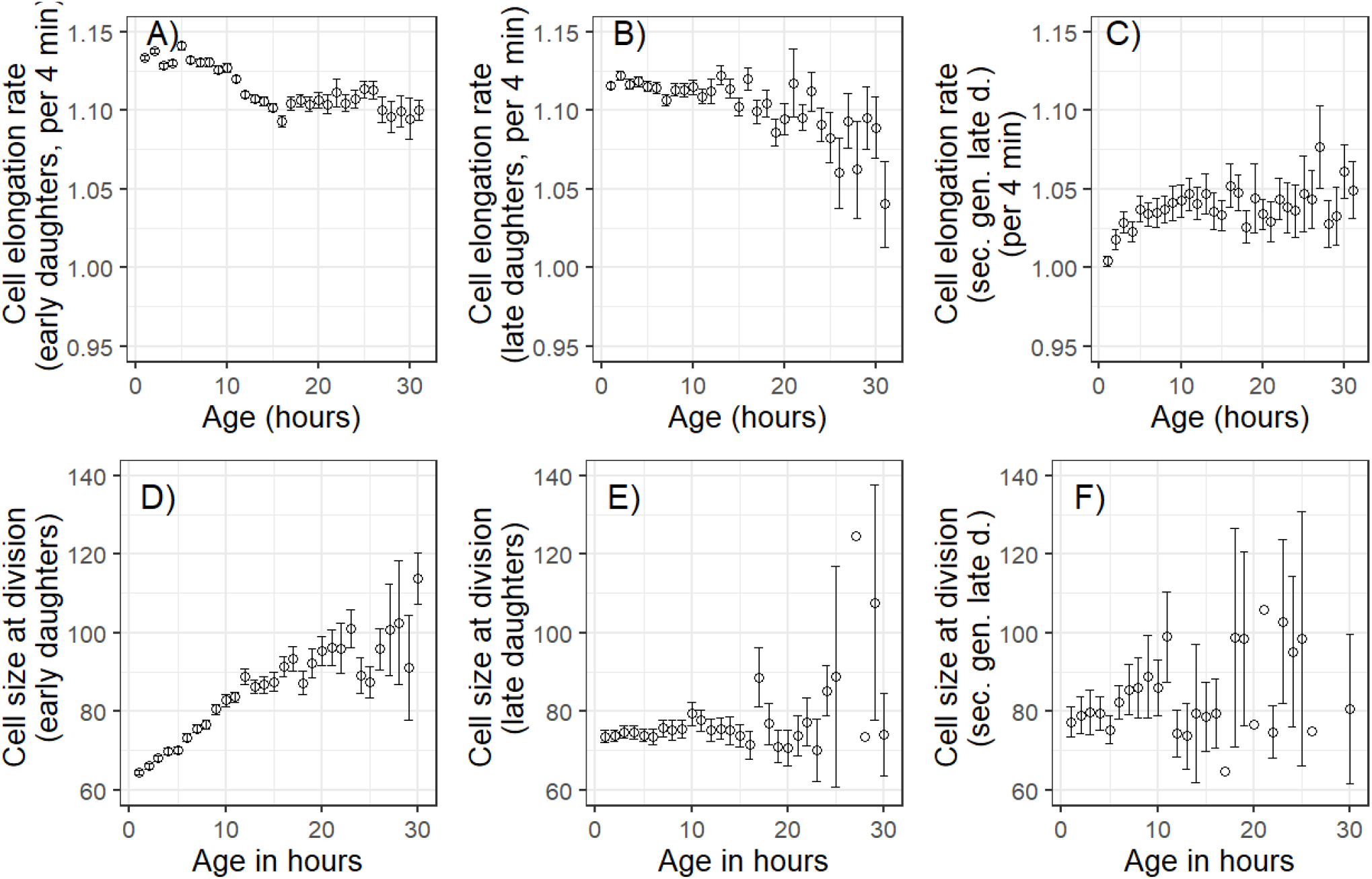
Cell elongation and cell size. A-C) Age-specific cell elongation rate (fractional elongation) and D-F) cell size at division (in pixels, 1pixel=0.064 μm) of the A&D) 516 early daughter cells, B&E) 516 late daughter cells, and C&F) 298 second generation late daughter cells. Hourly means ± standard errors are plotted. Associated model comparisons are presented in Table S1).

**Table S1:**
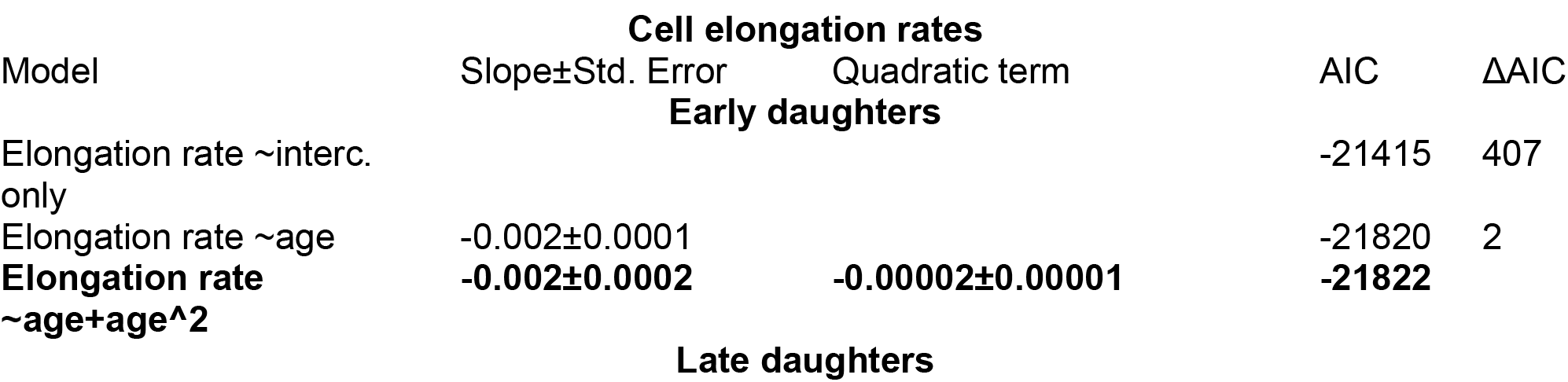
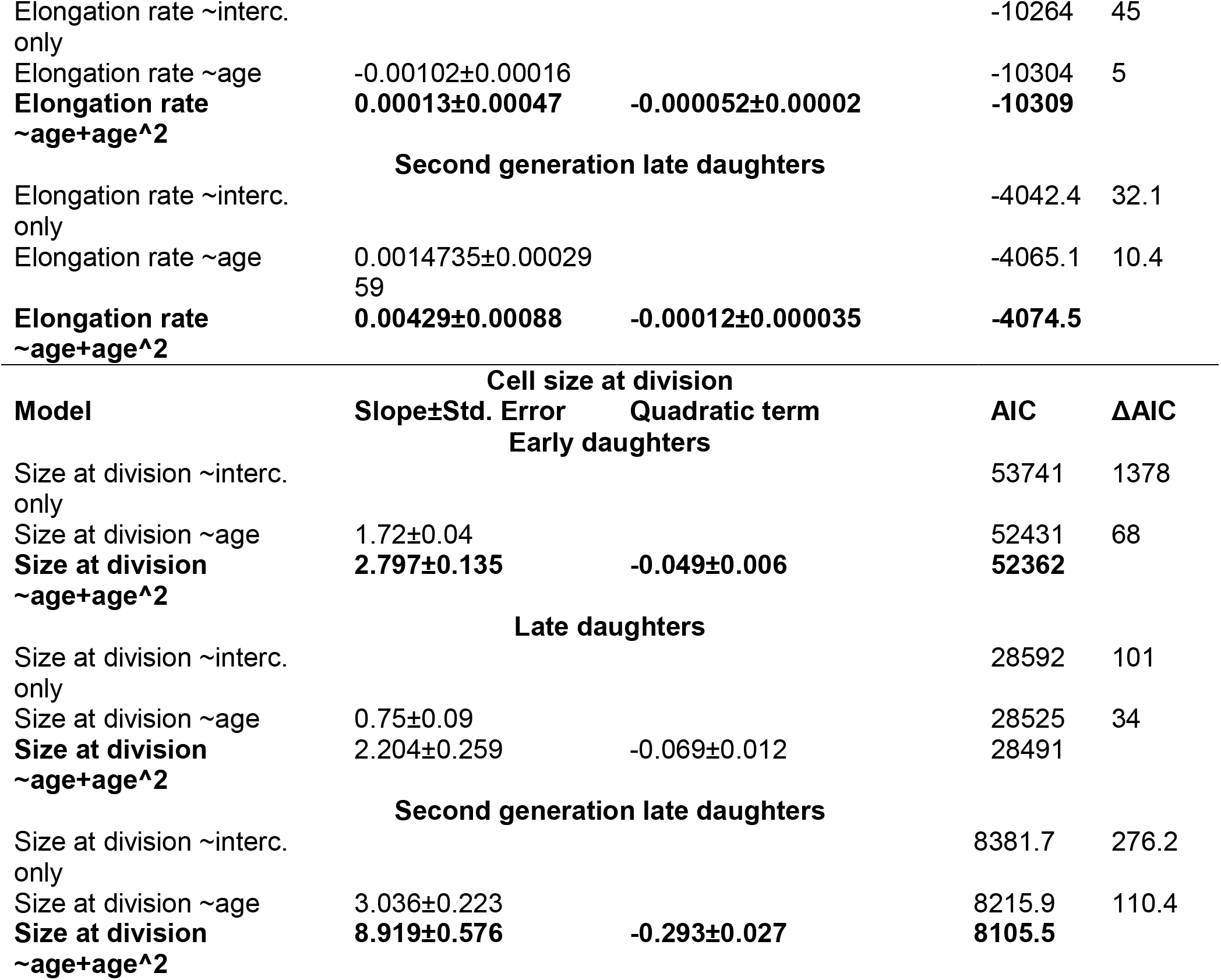
Model selection for age-specific cell elongation rates and cell size at division.

### Size at division

We illustrate cell size at division for early, late, and second generation late daughters (Fig. S2). When comparing linear models, the best fitted model for all three cell types was a model with a linear and quadratic term, though the curvature was minimal for early and late daughters (Table S1). Early daughters showed clear senescence in cell elongation rate, late daughters showed weaker senescence in cell elongation rate, and patterns for second-generation late daughters are less clear.

### Reverse time analyses: Hours before death

Chronological aging follows individuals from birth to death and time counted starts at birth. Various hypotheses about the evolution of life histories, aging and senescence, in particular, terminal investment strategies (Williams 1966, Clutton-Brock 1984, Charlesworth 1994), assume that individuals can sense that they are approaching death and that an optimal strategy for an individual might be to invest remaining resources into reproduction rather than maintenance when approaching immediate death. Such terminal investment has also been labeled as terminal illness, last year effect, or similar terms, and have been studied in a broad range of taxa, also using reverse time analyses (Coulson and Fairweather 2001). As Fig. 2 & Fig. 3 in the main text illustrates, age at death is highly variable, for that chronological senescence patterns and terminal investment strategies might provide very different insights. We investigated whether patterns in reproduction (division rate), cell elongation rate, or size at division differ significantly when time is counted backward (remaining lifespan), starting with death. We would expect that the last hour before death would show significant changes compared to the more spread out chronological senescence patterns (Fig. 2, Fig. S3 & S4). In general, there seems to be little evidence that terminal investment is happening to a larger degree, at least such strategies if existent, do not show larger effects compared to chronological aging and classical senescence (Fig. S5 & S6, Table S4 & S5).

### Division rate for hours before death

When individual cells approach death, their division rates decrease (Fig. S5), and these senescence patterns are similar compared to the (chronological) senescence patterns (Fig. 2A, B) (Table S4 & Table S2).

**Fig. S5:**
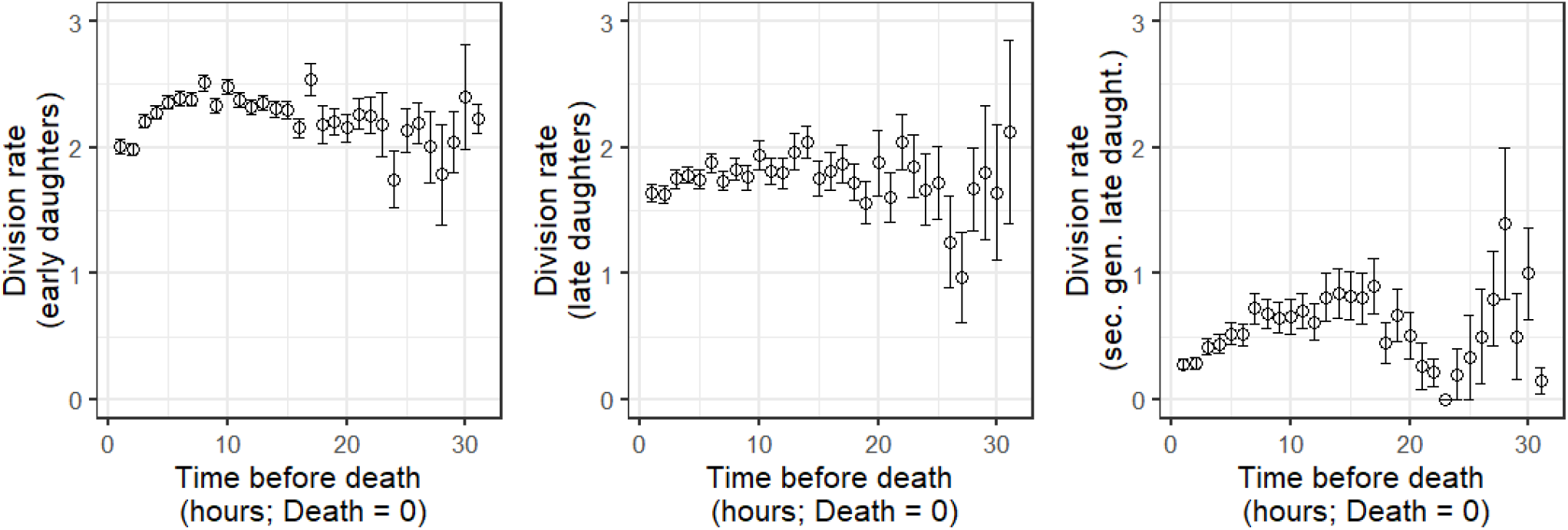
Hourly division rate for hours before death [remaining life time] (death = time 0) for left panel the 516 early daughters cells, mid panel the 516 late daughters, and right panel the 298 second generation daughters. Means ± standard errors are plotted. Associated model selection is shown in Table S4.

**Table S4:**
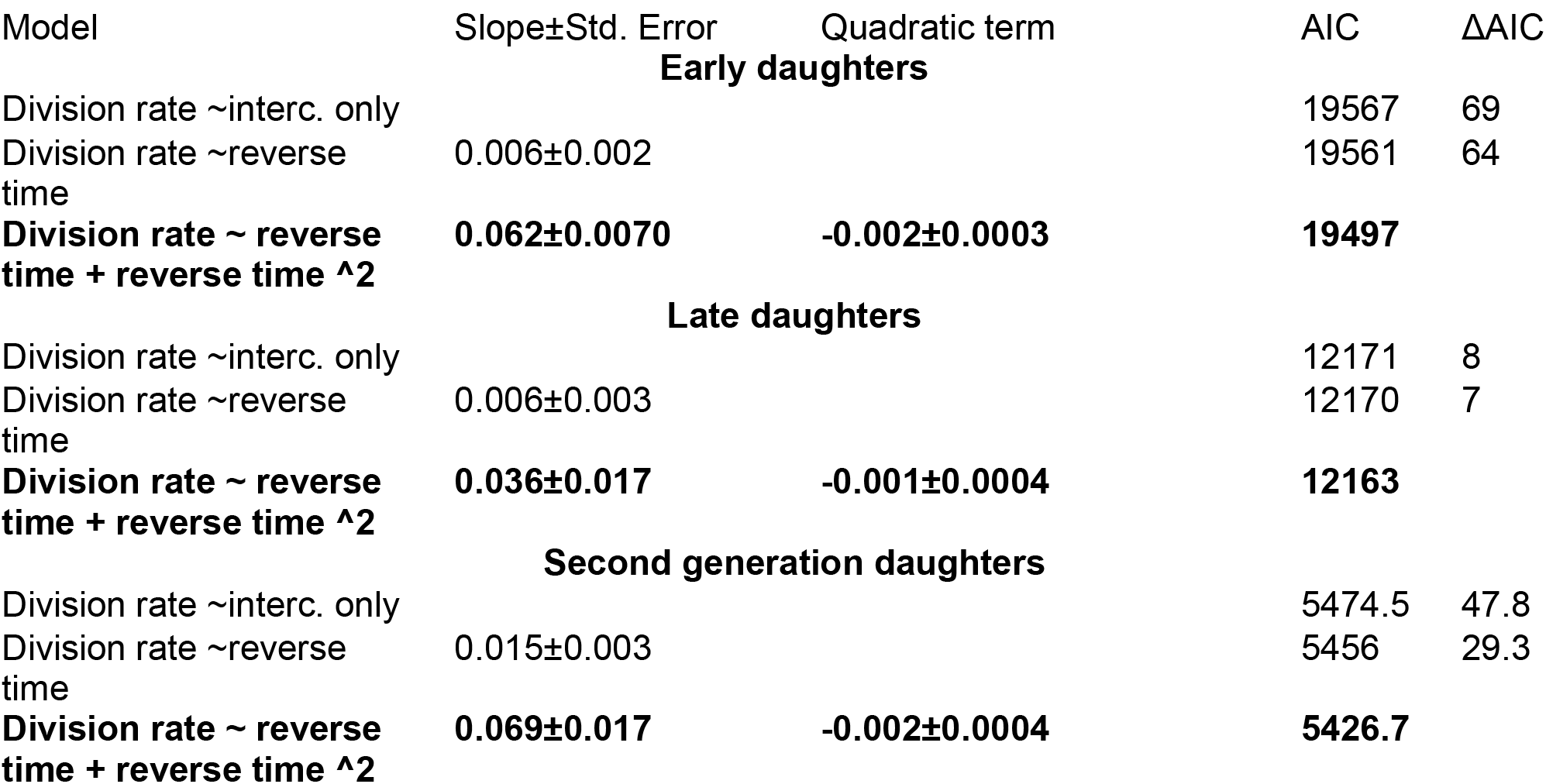
Model selection for division rate in reverse time (starting with age at death=0).

|  |  |  |  |  |
| --- | --- | --- | --- | --- |
| Model | Slope±Std. Error | Quadratic term | AIC | ΔAIC |
| Early daughters |  |  |  |  |
| Division rate ~interc. only |  |  | 19567 | 69 |
| Division rate ~reverse time | 0.006±0.002 |  | 19561 | 64 |
| **Division rate ~ reverse time + reverse time ^2** | **0.062±0.0070** | **-0.002±0.0003** | **19497** |  |
| Late daughters |  |  |  |  |
| Division rate ~interc. only |  |  | 12171 | 8 |
| Division rate ~reverse time | 0.006±0.003 |  | 12170 | 7 |
| **Division rate ~ reverse time + reverse time ^2** | **0.036±0.017** | **-0.001±0.0004** | **12163** |  |
| Second generation daughters |  |  |  |  |
| Division rate ~interc. only |  |  | 5474.5 | 47.8 |
| Division rate ~reverse time | 0.015±0.003 |  | 5456 | 29.3 |
| **Division rate ~ reverse time + reverse time ^2** | **0.069±0.017** | **-0.002±0.0004** | **5426.7** |  |

### Cell elongation rate for hours before death

The cell elongation rate does not drastically fall off when approaching death (no initial steep increase in cell elongation rate close to the time of death) (Fig. S6, Table S5). The senescence pattern is, at least for early and late daughters, less pronounced compared to that observed for chronological senescence (Fig. S2). Second generation daughters seem to reduce cell growth rates before death more substantially, but this reduction already starts a few hours before death and is not in line with terminal investment theories.

**Fig. S6:**
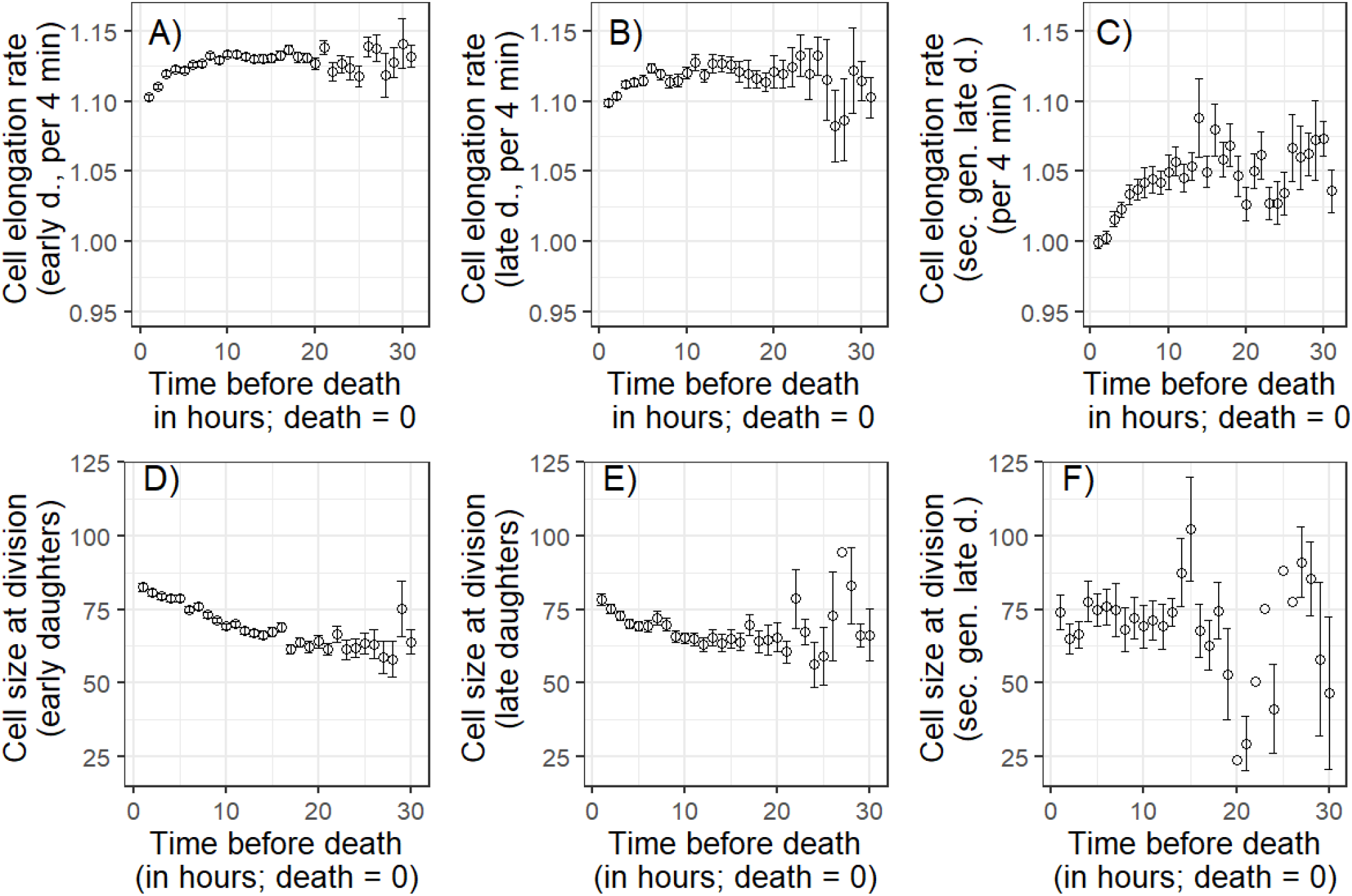
(Fractional) Cell elongation rate and cell size at division (in pixels, 1pixel=0.064 μm) for hours before death (death = time 0) for A & D) the 516 early daughters, B & E) the 516 late daughters, and C & F) the 298 second generation late daughters. Means ± standard errors are plotted. Associated model selection is shown in Table S5.

**Table S5:** Model selection for cell elongation and cell size at division rate in reverse time (starting with age at death=0).

| Cell elongation (reverse time) |  |  |  |  |
| --- | --- | --- | --- | --- |
| Model | Slope±Std. Error | Quadratic term | AIC | ΔAIC |
| Early daughters |  |  |  |  |
| Elongation rate ~interc. only |  |  | -22138 | 250 |
| Elongation rate ~reverse time | 0.001±0.00009 |  | -22288 | 100 |
| **Elongation rate ~ reverse time + reverse time ^2** | **0.004±0.0003** | **-0.0001±0.00001** | **-22388** |  |
| Late daughters |  |  |  |  |
| Elongation rate ~interc. only |  |  | -10529 | 65 |
| Elongation rate ~reverse time | 0.001±0.0001 |  | -10561 | 33 |
| **Elongation rate ~ reverse time + reverse time ^2** | **0.003±0.0004** | **-0.0001±0.00002** | **-10594** |  |
| Second generation late daughters |  |  |  |  |
| Elongation rate ~interc. only |  |  | -4082.8 | 122.1 |
| Elongation rate ~reverse time | 0.003±0.0003 |  | -4167.5 | 37.4 |
| **Elongation rate ~ reverse time + reverse time ^2** | **0.008±0.0009** | **-0.0002±0.00003** | **-4204.9** |  |
| Cell size at division (reverse time) |  |  |  |  |
| Early daughters |  |  |  |  |
| Size at division ~interc. only |  |  | 53679 | 386 |
| Size at division ~reverse time | -0.952±0.0486 |  | 53299 | 6 |
| **Size at division ~ reverse time + reverse time ^2** | **-1.337±0.145** | **0.018±0.006** | **53293** |  |
| Late daughters |  |  |  |  |
| Size at division ~interc. only |  |  | 28629 | 26 |
| Size at division ~reverse time | -0.438±0.088 |  | 28606 | 3 |
| **Size at division ~ reverse time + reverse time ^2** | **-0.990±0.258** | **0.026±0.011** | **28603** |  |
| Second generation late daughters |  |  |  |  |
| Size at division ~interc. only |  |  | 8182.3 | 54.9 |
| Size at division ~reverse time | 1.254±0.236 |  | 8156.6 | 29.2 |
| **Size at division ~ reverse time + reverse time ^2** | **4.531±0.627** | **-0.158±0.028** | **8127.4** |  |

### Cell size at division for hours before death

Cell size at division increased with chronological age, but mainly for early daughters (Fig. S2). One mechanism that might lead to such an increase in cell size at division is the formation of filamentous cells: cells that do continue to elongate but stop dividing. Such filamentation is usually seen as a stress response and increases mortality of the cell (Mattick et al. 2003, Wang et al. 2010). Previous studies (conducted under different media and temperature than ours) have reported higher filamentation rates with increasing age (Wang et al. 2010). Under our growth conditions, cells that entered a filamentation stage rarely recovered from that stage and did not return to a regular dividing smaller cell, though they not always stalled dividing. To test if such filamentous cells drive the chronological senescence pattern in size at division, we investigated the cell size for the hours before death. For early and late daughters, we detected an increase in cell size at division before death, but this increase is already initiated hours before the actual death and not a final increase in size due to filamentation (Fig. S6, Table S5).

### Random deterioration model

Our random deterioration model builds on a LeBras type cascading failure model(Le Bras 1976). In such discrete state models, individuals increase from a current state *i* to a state *i+1* at a rate proportional to i, and mortality increases proportionally to state i. We can think of such states as accumulating damage, or model the reverse process, where an individual starts with some vitality and this vitality then decreases with increasing state *i* (random deterioration) (Weitz and Fraser 2001). In this study, we assume accumulating damage that stands symbolically for an unknown aging factor. These random deterioration models are mathematically closely linked to demographic frailty models that do not assume a stochastic stage progression among individuals, but rather heterogeneity of individual is defined by fixed frailty at birth (Yashin et al. 1994). Such frailty, an age independent mortality that follows a gamma distribution (Missov and Vaupel 2015), can be added to the classical models in demography, a Gompertz- model, which describes an age-dependent exponential increase in mortality. These models can also be extended by a Makeham term, an age-independent baseline mortality, which further improves model fit, at least for humans and various other complex species.

We first estimated parameters of such a Gamma-Gompertz-Makeham (GGM) model, *Zae^bx^*+*c* fitted to the survival data of the early daughters. *Z* is the random frailty variable that follows a gamma distribution across individuals. The distribution has a mean 1 and variance of σ^2^. This definition of a gamma distribution with a mean of 1 implies that we have to set the scale parameter equal to the shape parameter of the gamma distribution. The rate of aging, *b*, is the same for all individuals, *x* determines the age in hours, *a* describes the baseline mortality of the Gompertz part of the model, and *c* describes the Makeham term, the age independent mortality.

We then transformed these estimates of the GGM model to a random deterioration model according to Yashin et al. (Yashin et al. 1994), making use of mathematical similarities between the GGM and random deterioration models.

Let *t(i)* = *λ*_0_+ *i λ*, *m*(*i*) = μ0 + *iμ* (for *i* greater than or equal to 0) denote the state transition rate and the death rate, respectively.

μ0=a+c, the death rate at birth i.e. state *i*=0,

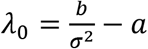, the transition rate from state *i*=0 to state *i*=1,

*λ* = *b* — *σ*^2^*a*, the change in the state transition rate between state *i* and *i*+*1*, for *i*>*0*

*μ* = *σ*^2^*a*, the change in the death rate at state *i*, for *i*>*0*

Resulting estimates for early daughters were μ0=0.056, *λ*_0_ = 0.178, *λ* = 0.598, and *μ* = 0.00023. Using these estimates for early daughters, we estimated the probability of observing an individual at state *i* at age *x*, *P*(*x,i*),

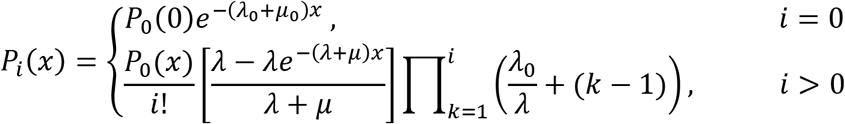

We then built a matrix of *P*(*x,i*) for which we limited the age, *x*, between 0 and 31 and the states were limited between 0 and 5000. The column sum of this matrix equals the survival probability at any given age *x* to age *x*+*1*.

### Microsimulations of the random deterioration model

In order to simulate the (damage) state at death distribution we used a microsimulation. As a step-by-step calculation would take too long, we modelled the state at death distribution by drawing of a Poisson distribution with a rate of 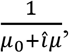, where 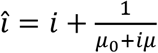. In doing so, we estimated the expected state at death, rather than tracking individuals transitioning through damage states and their associated mortality risk as a step-by-step calculation would do. A step-by-step simulation would take a long time because individuals can easily end up in very high damage states (very long tailed distributions).

### Estimating parameters for the late daughters

For the late daughters, we used the same state transition and survival parameters (μ0=0.056, *λ*_0_ = 0.178, *λ* = 0.598, and *μ* = 0.00023) that we estimated for the early daughters. We did so because early and late daughter cells should biologically not be fundamentally different – aside of starting with different damage levels at birth. They should accumulate damage at the same rate and should experience mortality risk just based on their damage state. We assumed that the late daughters are born at the same stage, or a scaled version of the same stage, in which the mothers died, therefore let

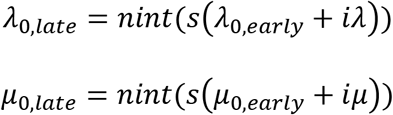

Where subscript *late* and *early* denotes late and early daughter estimates and *s* ∊ (0,1) and *nint*(.) is the nearest integer function. So based on the estimated damage state distribution of the mothers at death (early daughters), we also knew the damage state distribution at birth of the daughters (late daughters) [see below for optimally scaling this distribution]. Using the state transition probabilities of the early daughters and the probabilities of death, we estimated the survival of the late daughters. We also estimated the survival functions of cells that did not change their damage state with age. Each grey line in main text Fig. 3, describes such a fixed damage class survival function.

In order to determine the transmission fraction of accumulated damage between mothers (early daughters) and daughters (late daughters), we optimized the scaling of the mother state at death distribution (fraction of mother-daughter transmission) to the daughter age at birth distribution. The optimization was based on observed survival of the late daughters, by minimizing the squared deviations from the observed survival function. This optimal scaling factor was estimated at 7%.

Similar to the late daughters, we simulated the damage state distribution at birth, the damage state distribution at death, and calculated the matrix *P*(*x*, *i*) for the second generation late daughters.

**Fig. S9:**
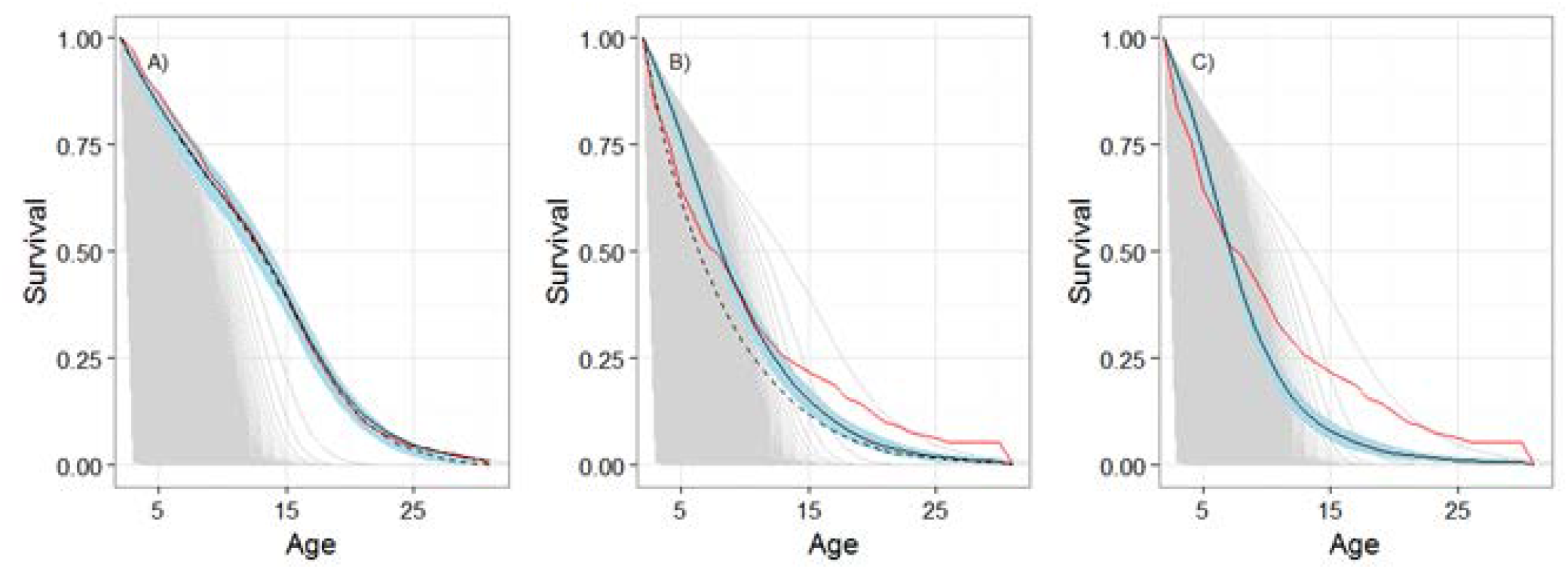
Observed population level survival curve (red line), GGM simulated survival curve with symmetric damage transmission (hatched black line) as well as GGM simulated survival curve with asymmetric damage transmission of 7% (solid black line ± 95% CI in blue) for early (A), late (B) and second generation late daughters (C). Graphs A & B are identical to Fig. 1C & D. Thin grey lines in depict expected survival curve of cells with different fixed damage state. That is, for instance, the outermost thin grey line in B depicts the survivorship curve of a hypothetical cohort that starts without damage and never accumulates any damage. The most left survival curve illustrates the low survival of cells that were born with maximum damage level of 5000.

**Fig. S10:**
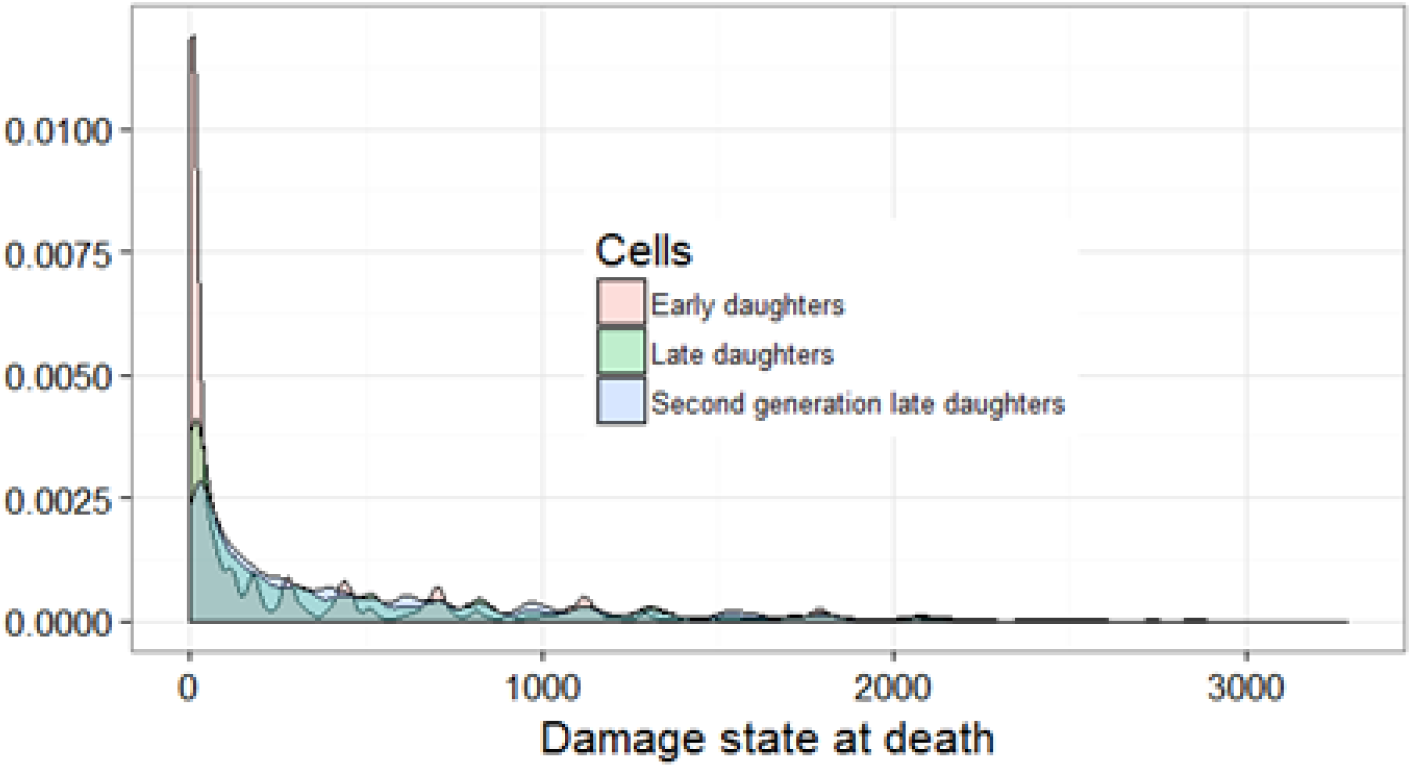
Density distribution of damage state at death of early daughters, late daughters, and second generation late daughters. Most cells die with little accumulated damage. Note, the not so smooth tails of the distributions are a result of drawing from a Poisson distribution in our simulations.

**Fig. S11:**
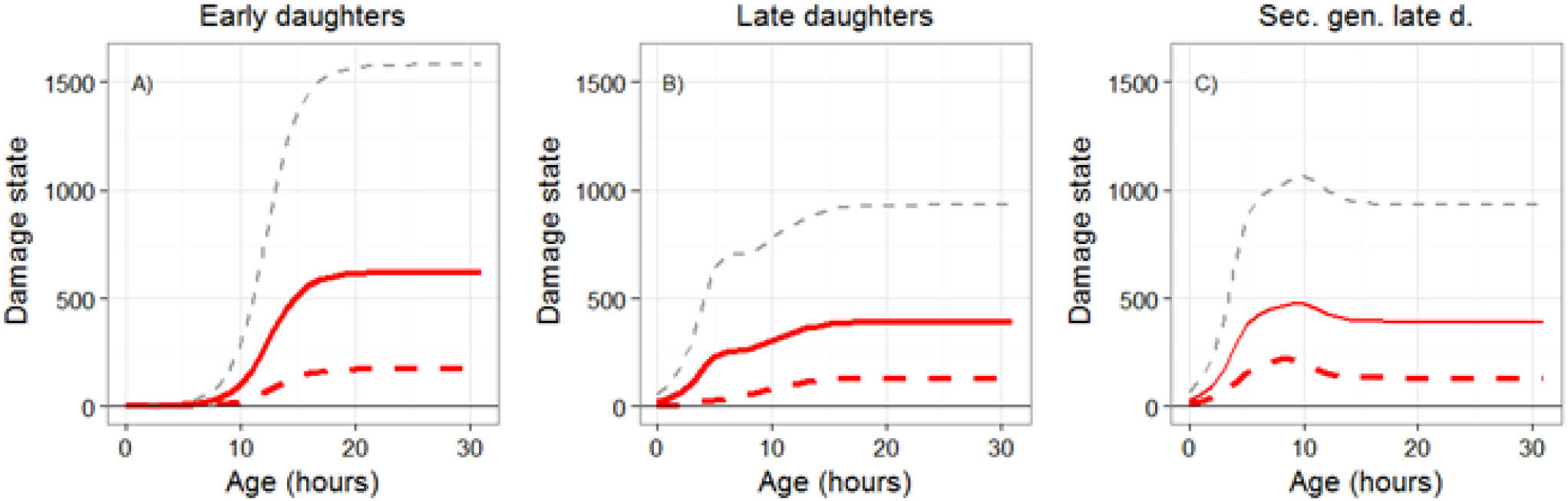
GGM model results: mean (red solid line, + SD grey hatched lines) and median (red hatched line) damage state with increasing age for early daughter cells (A), and late daughter cells (B) and second generation late daughter cells (C). Panel A & B are identical to Panel Fig. 1A, B.

**Fig. S12:**
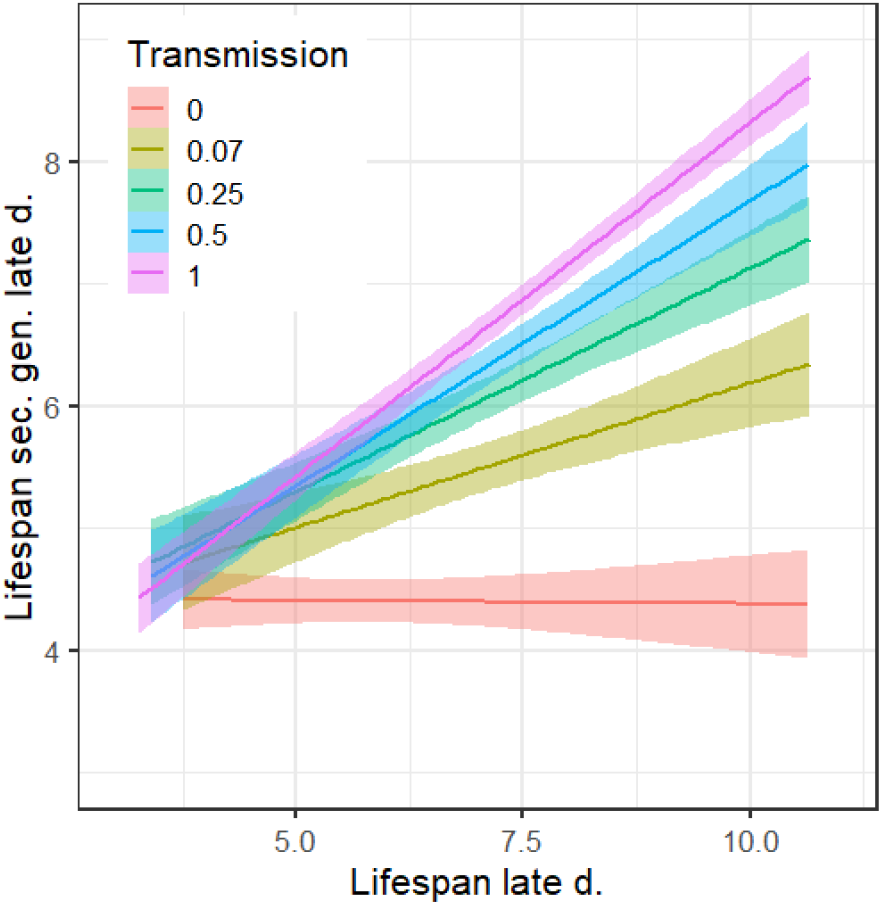
Lifespan (square root transformed) of simulated 516 late daughter cells (mothers) versus the lifespan of their simulated last daughter cell (second generation late daughter cells) with different levels of mother to daughter damage transmission. Scenarios include the optimized fixed transmission level at 0.07 (forest green), a scenario for perfect rejuventation, i.e. 0 transmission (red), 0.25 transmission (green), 0.5 transmission, i.e. symmetric (equal) transmission between mother and daughter (blue), and transmission of all accumulated damage to the daughter (1) (pink).CI are shown for each correlation in lifespan between mothers and daughters as shaded areas.

Authors contributions: UKS designed the study, UKS, MN & PC performed the experiments, UKS, AL, XS analyzed the data, all authors substantially contributed to discussions and writing the manuscript, UKS wrote the first and final draft of the manuscript.

